# Ablation of hematopoietic stem cell derived adipocytes reduces tumor burden in syngeneic mouse models of high-grade serous carcinoma

**DOI:** 10.1101/2024.09.19.613924

**Authors:** Elizabeth R Woodruff, Courtney A Bailey, Francis To, Vyshnavi Manda, Joanne K Maltzahn, Timothy M Sullivan, Meher P Boorgula, Maria Sol Recouvreux, Ruby Vianzon, Bogi Conrad, Kathleen M Gavin, Kimberly R Jordan, Dwight J Klemm, Sandra Orsulic, Benjamin G Bitler, Zachary L Watson

## Abstract

Hematopoietic stem cell-derived adipocytes (HSCDAs) are an adipocyte subtype derived from myeloid precursors. We hypothesized that HSCDAs promote high-grade serous ovarian carcinoma (HGSC). Despite similar adipogenic differentiation, primary human female HSCDAs exhibit transcriptional differences from conventional mesenchymal adipocytes (CMAs), including upregulation of metabolic pathways. HSCDAs secreted greater amounts of IL-6 and IL-8 than CMAs. We generated immunocompetent HSCDA Proficient (HSCDA-Pro; can make both adipocyte subtypes) and Deficient (HSCDA-Def; can only make CMAs) mice and conducted two independent tumor studies using two unique syngeneic HGSC models. Tumor burden trended lower in HSCDA-Def mice in both models. Omentum tumors from HSCDA-Def mice exhibited downregulated transcription of metabolic pathways. Tumors from HSCDA-Def mice also had lower densities of dendritic cells (DC) and natural killer (NK) cells. These data suggest that HSCDAs promote HGSC progression through metabolic and immune pathways.

## INTRODUCTION

Most ovarian cancers are of the high-grade serous subtype (high-grade serous carcinoma; HGSC) [1]. HGSC preferentially invades visceral adipose tissues (VAT) such as the omentum [2–5], a complex apron-like fold of the visceral peritoneum that lies across the anterior aspect of the intestines. It is primarily composed of adipocytes surrounded by a mesothelial layer but also contains vasculature and “milky spots,” small tertiary lymphoid aggregates that may be innervated by sympathetic nervous system fibers [6–8]. Omental tumors are often the largest abdominal tumors in women with ovarian cancer, suggesting that the microenvironment of the omentum is highly conducive to HGSC cell invasion and proliferation. Adipocytes actively enhance HGSC metastasis through several mechanisms, including but not limited to: (a) secretion of inflammatory cytokines, such as interleukin-8 (IL-8), promoting HGSC migration and invasion, (b) providing energy to HGSC cells through direct lipid transfer, and (c) stimulating upregulation of fatty acid-binding protein 4 (FABP4), fatty acid translocase (CD36), and other regulatory proteins of fatty acid transport and lipid metabolism [9–11].

During aging, visceral adiposity increases through the enlargement of existing adipocytes (hypertrophy) and/or *de novo* production of new adipocytes (hyperplasia). Adipocytes were previously proposed to arise exclusively from mesenchymal precursors; however, a 2006 murine bone marrow transplant study demonstrated that a subset of circulating myeloid cells home to adipose tissue, become resident, and terminally differentiate into adipocytes [12]. These cells were termed hematopoietic stem cell-derived adipocytes (HSCDAs) [13] to distinguish them from conventional mesenchymal-derived adipocytes (CMAs). Published studies confirmed the presence of HSCDAs in humans [14, 15]. Ryden et al. isolated mature white adipocytes from recipients of bone marrow transplants (BMT) and evaluated them for donor and recipient DNA. Up to 27% of purified adipocytes were derived from donor cells and the fraction of donor-derived adipocytes was positively correlated with time since transplant [14]. Gavin et al. found that up to 35% of mature adipocytes were generated from donor cells in BMT recipients, and the proportion of donor DNA was positively correlated with elapsed time post-transplant [15]. These two independent studies both confirmed human HSCDA existence and demonstrated that their proportion of total adipose cells increases with age.

Subsequent studies developed methods to isolate and differentiate human primary adipocyte precursors into HSCDA and CMA populations and established an innovative murine model for HSCDA ablation [13, 16–18]. These studies demonstrated that mouse HSCDAs accumulated with age and occurred in significantly higher numbers in female versus male mice. Transcription analysis showed that HSCDAs differ from CMAs by upregulation of inflammatory gene expression and decreased expression of leptin and lipid oxidation pathways, among others [19]. Of key relevance to menopause and HGSC, mouse HSCDA production was found to be enhanced by models of “pseudomenopause” such as ovariectomy and estrogen receptor alpha knockout. HSCDA production in ovariectomized mice was attenuated by estradiol replacement, demonstrating that the ovarian hormone is necessary and sufficient for the constraint of HSCDA production. In agreement with prior anatomical studies, HSCDA accumulation in these mice was higher in VAT and gonadal adipose tissue (GAT) compared to subcutaneous adipose tissue (SAT) depots [17]. Finally, depletion of HSCDAs in mice did not alter the quantities of tissue-resident immune cells in GAT or SAT depots; however, HSCDA-depleted mice were found to have increased circulating leptin levels, indicating that the presence or absence of HSCDAs may alter lipid homeostasis [13].

Given their presence in humans and their characteristics in murine models, we initiated studies of HSCDAs in the context of ovarian cancer, including comparisons of HSCDA and CMA transcription and cytokine secretion, as well as tumor studies in HSCDA ablated mice.

## RESULTS

### Human hematopoietic and mesenchymal precursor cells are equally capable of adipogenic differentiation

Studies have successfully differentiated mouse hematopoietic and mesenchymal precursor cells into HSCDAs and CMAs, respectively [13]. To develop a complementary human *in vitro* model, researchers collected human subcutaneous adipose biopsies, which were dissociated, sorted by fluorescence-activated cell sorting (FACS) into mesenchymal and hematopoietic populations, and then cryopreserved for subsequent adipogenic differentiation. For this study, with informed consent for secondary use, we acquired adipose progenitors from human female donors. As the original human subjects research that collected these tissues is in the preprint stage [18], with permission from the authors, we have included information about study participants, adipose biopsy collection and processing, flow sorting, and subsequent culture in our Materials and Methods. The FACS gating strategy is shown in **Supplementary Fig. S1**. Study participant data, including age, menopause status, height, total mass, and fat mass are shown in **Table 1**.

**Table 1:** Adipose Donor Characteristics.

| Donor ID | Age | Menopause | Height (cm) | Fat mass (kg) | Total mass (kg) | Relative fat mass (%) |
| --- | --- | --- | --- | --- | --- | --- |
| 1 | 52 | Post | 173 | 32 | 79 | 41 |
| 2 | 53 | Post | 180 | 41 | 90 | 46 |
| 3 | 56 | Post | 157 | 16 | 50 | 31 |
| 4 | 33 | Pre | 165 | 33 | 83 | 40 |
| 5 | 26 | Pre | 157 | 30 | 77 | 39 |
| 6 | 54 | Post | 170 | 20 | 58 | 34 |
| 7 | 52 | Post | 160 | 29 | 73 | 40 |
| 8 | 53 | Post | 168 | 50 | 111 | 45 |
| 9 | 61 | Post | 173 | 34 | 83 | 41 |

To determine if human mesenchymal and hematopoietic progenitor cells have a similar capacity for adipogenic differentiation, we acquired daily brightfield images of differentiating cells from three independent female postmenopausal human donors (Donors 1-3). **Fig. 1A-C** shows cells from each donor at the pre-differentiated state (day 0, D0), mid-differentiation (day 8, D8), and at the end of the differentiation process (day 14, D14). We grew precursor cells to confluence. Notably, cells from Donor 2 did not pack as densely as those from Donors 1 and 3 (D0). By D8, all cells began to show accumulation of lipid droplets, which were extensive by D14 in both Donor 1 and Donor 3 **(Fig. 1A, C)**. Both mesenchymal and hematopoietic cells from Donor 2 showed less extensive lipid accumulation **(Fig. 1B)**. The full set of daily images (D0-D14) for all three donors are shown in **Supplementary Fig. S2-S4**.

**Fig. 1.**
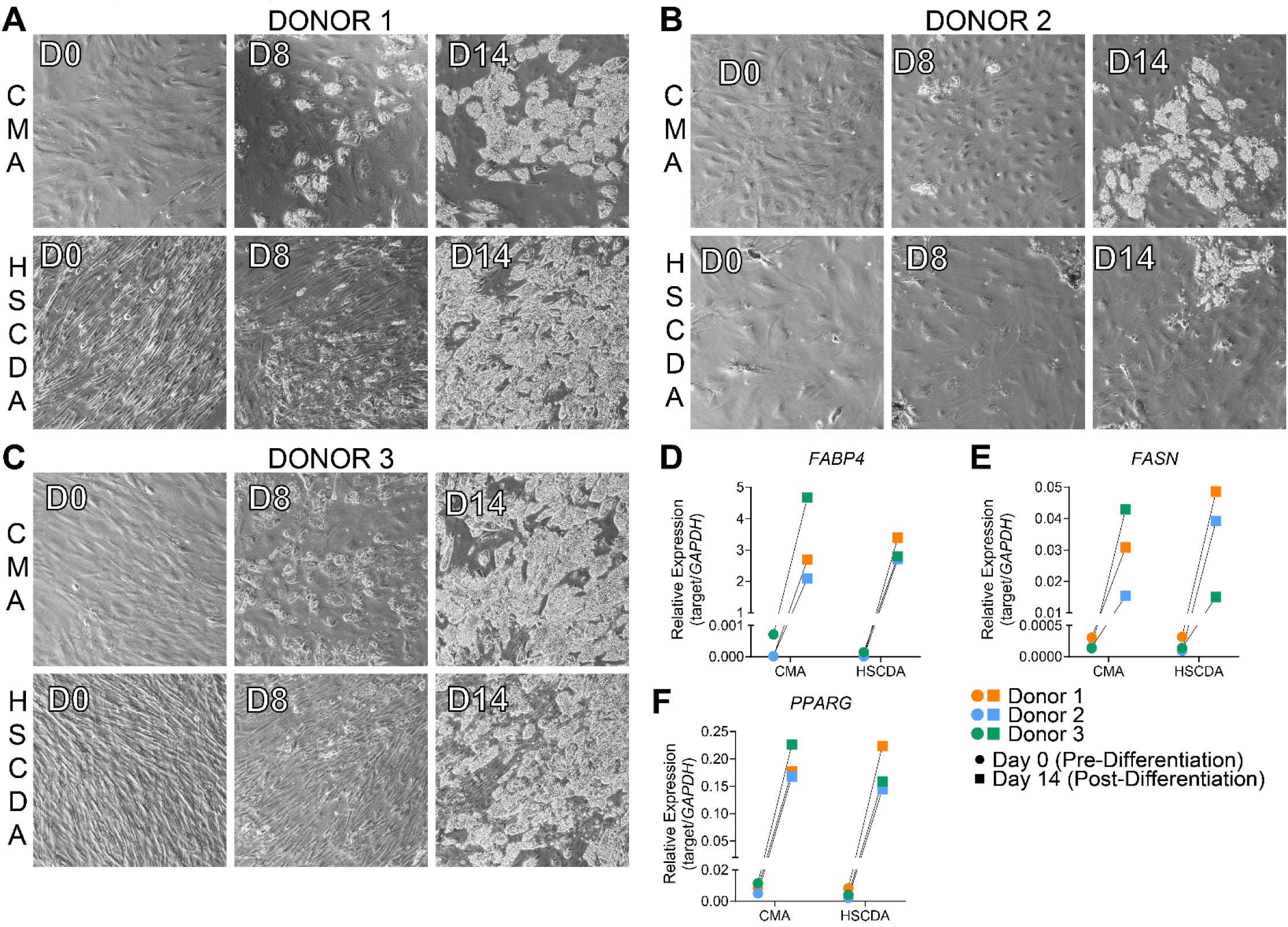
Human hematopoietic and mesenchymal precursors undergo similar adipogenic differentiation *in vitro*. **(A-C)** Brightfield images of cells from human female Donors 1-3 as they progress from confluent pre-differentiated precursor cells (D0) through mid-differentiation and appearance of lipid droplets (D8) to post-differentiation (D14). Paired mesenchymal precursor/CMAs and hematopoietic/HSCDAs are shown. **(D-F)** Confluent human precursor cells and fully differentiated HSCDAs and CMAs were analyzed by RT-qPCR for mRNA expression of *FABP4*, *FASN*, and *PPARG*. Each data point is the mean of triplicate reactions, and expression is normalized to *GAPDH*. Paired data points show relative expression for cells from each donor at D0 (circles) and D14 (squares) .

To further compare adipogenic differentiation, we isolated RNA from donor cells collected pre-differentiation (D0) and post-differentiation (D14) and performed RT-qPCR to determine if adipose gene expression (*FABP4*, *FASN*, and *PPARG*) was similar between HSCDAs and CMAs. All cells from all three donors showed similar ranges of relative target gene expression at D0 and higher expression at D14 **(Fig. 1D-F)**. For example, relative *PPARG* expression (compared to *GAPDH*) increased by an average of 34.1-fold in CMAs, and 42.9-fold in HSCDAs, with no significant differences (paired t-test; p=0.1735). Comparable results were observed in two biological replicates **(Supplementary Fig. S5)**, indicating that myeloid precursor cells are capable of similar adipogenic differentiation to mesenchymal cells.

### Cultured human HSCDAs and CMAs exhibit differential transcription

Prior microarray transcriptomic data from murine-derived HSCDAs and CMAs showed that the former exhibit a highly inflammatory transcriptional profile compared to the latter, including increased mRNA expression of cytokines [19]. We hypothesized that *in vitro* cultured human HSCDAs would also exhibit unique transcriptional profiles. We performed RNA-Seq on matched HSCDA/CMA pairs (post-differentiation at D14) from three independent donors (Donors 1-3) **(Fig. 2A)**. Principal component analysis (PCA) showed that HSCDA and CMA cells from Donor 1 and Donor 3 clustered together, while HSCDAs and CMAs did not cluster for Donor 2 **(Fig. 2B)**. Despite these differences, and the less extensive lipid accumulation in cells from Donor 2 **(Fig. 1)**, we used data from all three donors and examined differential gene expression (DGE, **Fig. 2C**) and performed over-representation analysis (ORA) for Hallmark pathways. Multiple cell cycle pathways decreased in human HSCDA cells, including E2F Targets, Mitotic Spindle, and G2M Checkpoint **(Fig. 2D)**. The heatmap for Hallmark E2F targets showed that HSCDAs, regardless of donor, clustered separately from CMAs **(Fig. 2E)**. Functional gene set enrichment analysis (fGSEA) recapitulated downregulation of these three pathways **(Fig. 2F)** and revealed enrichment in HSCDAs for multiple Hallmark metabolic pathways, including Oxidative Phosphorylation, Adipogenesis, and Fatty Acid Metabolism **(Fig. 2F-I)**. Overall, these data show that, despite variability in gene expression and differentiation from specific donors, HSCDAs exhibit a unique transcriptional profile from CMAs, including differences in cell cycle and metabolic pathways.

**Fig. 2.**
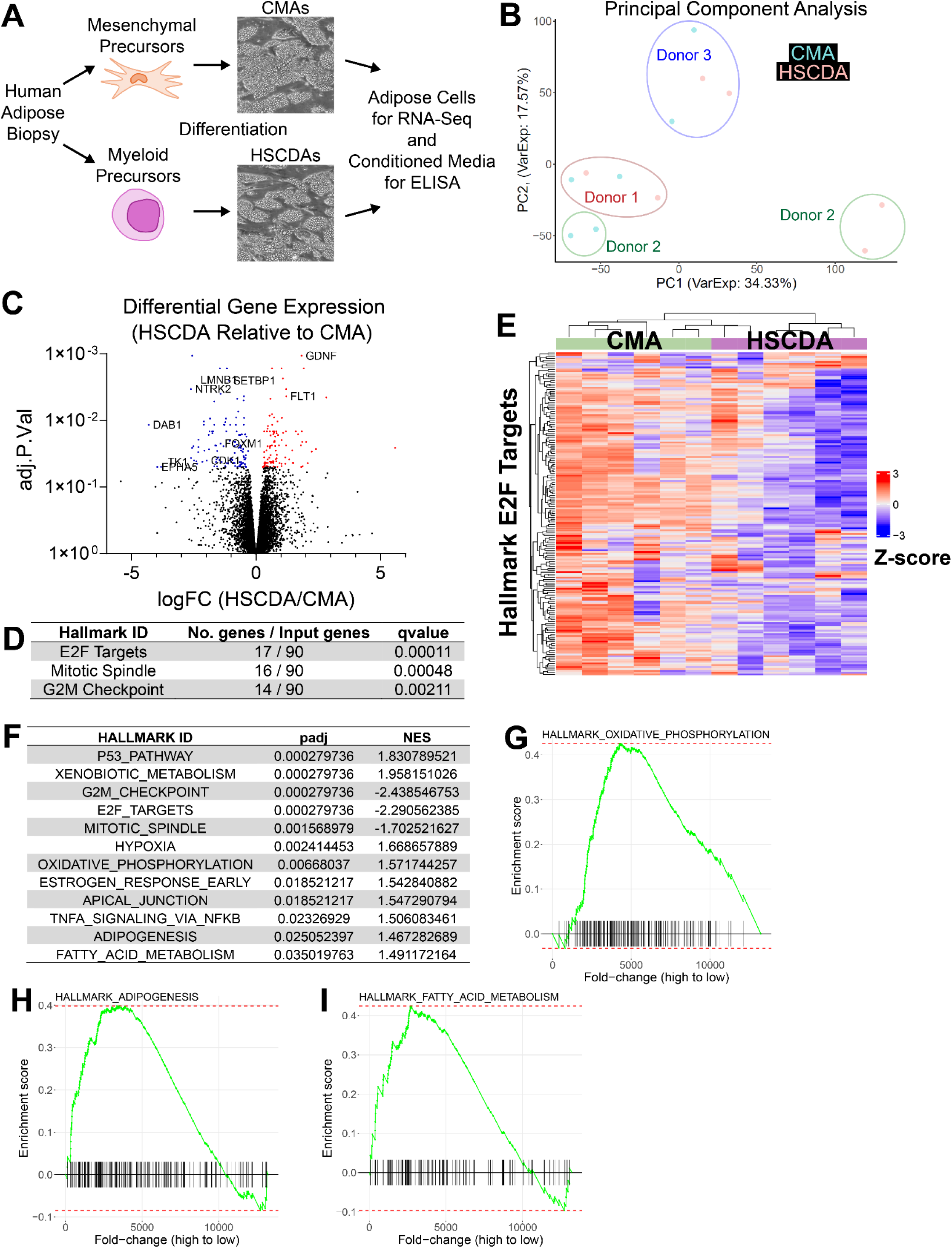
*In vitro* differentiated human primary HSCDAs exhibit unique transcription relative to CMAs. **(A)** Diagram of adipogenic differentiation of human primary precursor cells into HSCDAs and CMAs. Cell transcriptomes were analyzed in duplicate by RNA-Seq. **(B)** Principal component analysis (PCA) of adipocyte RNA-Seq. **(C)** Volcano plot of differential gene expression (DGE) comparing HSCDAs relative to CMAs. Specific gene names in E2F and other pathways are highlighted. **(D)** Significantly differentially regulated hallmark pathways by over-representation analysis (ORA). **(E)** Heatmap of Hallmark E2F Targets showing transcriptional downregulation of the pathway in HSCDAs relative to CMAs. **(F)** Hallmark pathways were significantly enriched in functional gene set enrichment analysis (fGSEA). padj = p value adjusted for multiple comparisons. NES = Normalized Enrichment Score. A positive NES indicates over-enrichment of pathway genes in HSCDAs relative to CMAs, while a negative NES indicates under-enrichment in HSCDAs relative to CMAs. Note the negative NES values for the hallmark pathways shown by ORA in **D**, confirming downregulation of these pathways in HSCDAs. **(G-I)** fGSEA plots corresponding to the indicated Hallmark metabolic pathways in **F**. The peak-shaped plots correspond to the positive NES and indicate over-enrichment in HSCDAs relative to CMAs.

### Cultured human HSCDAs secrete greater amounts of inflammatory cytokines

While we did not observe transcriptional differences in inflammatory cytokines in human HSCDAs that were seen in mouse HSCDAs [19], we hypothesized that secretion may differ between human HSCDAs and CMAs. We analyzed conditioned media from paired HSCDAs and CMAs obtained from six additional independent human female donors (Donors 4-9) using a multiplex ELISA for 10 cytokines (IFN-γ, IL-1β, IL-2, IL-4, IL-6, IL-8, IL-10, IL-12p70, IL-13, and TNF-α). HSCDAs exhibited increased IL-6 and IL-8 secretion, with four donors producing higher concentrations of IL-6 and five donors producing higher concentrations of IL-8 compared to CMAs **(Fig. 3A-F)**. Averaged across all six donors, HSCDA media contained 5.7-times the concentration of IL-6, and 6.7-times the concentration of IL-8, compared to paired CMA media. The remaining cytokines were not detected within the range of the assay.

**Fig. 3.**
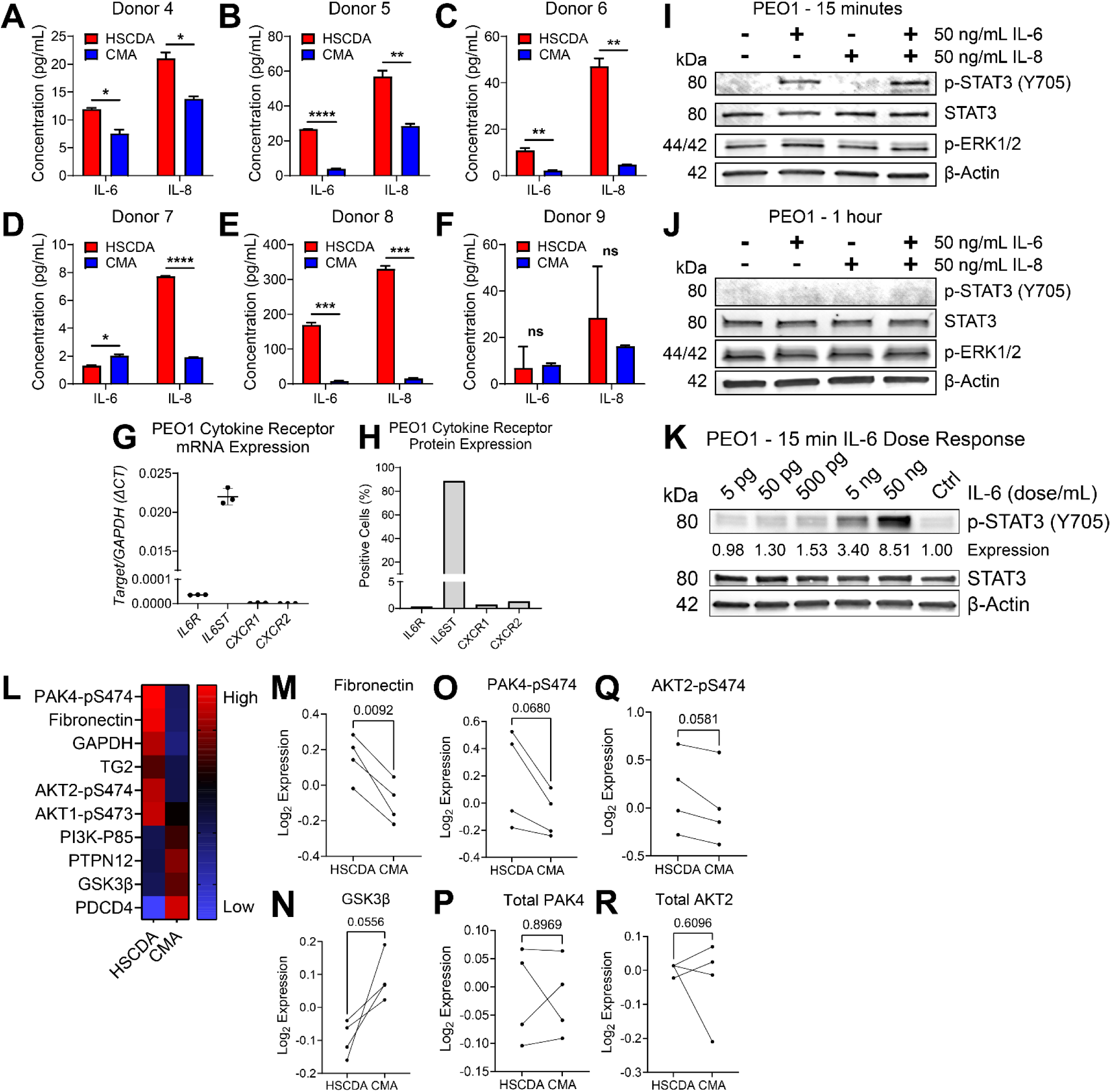
*In vitro* differentiated human primary HSCDAs secrete greater amounts of IL-6 and IL-8 than CMAs. **(A-F)** HSCDA or CMA conditioned media from six independent human donors was examined by ELISA for IL-6 and IL-8. Data are plotted as mean ± SD of two wells. Unpaired t-test. * p<0.05, ** p<0.01, *** p<0.001, **** p<0.0001. **(G)** RNA was isolated from PEO1 cells and RT-qPCR was performed for the indicated cytokine receptor transcripts. *GAPDH* was used for ΔCt quantification. **(H)** Flow cytometry was performed on PEO1 cells to detect surface expression of the indicated cytokine receptor proteins. **(I-K)** PEO1 cells were incubated with the indicated cytokine supplements for the indicated time. Immunoblots were performed for phosphorylated and total STAT3 (IL-6 signaling), phosphorylated ERK1/2 (IL-8 signaling), and β-actin loading control. **(L-R)** PEO1 cells were incubated in conditioned media from HSCDAs or CMAs for 24 hours. Protein expression and phosphorylation were examined by RPPA. **(L)** The heat map shows the average median centered Log2 expression for PEO1 cells incubated in 4 independent media pairs (Donors 6 through 9). **(M-N)** Representative paired scatter plots showing consistent differential protein expression of fibronectin and GSK3β in PEO1 cells incubated in each media pair. **(O-R)** Paired scatter plots for phosphorylated and total forms of PAK4 and AKT2, showing differential regulation of serine phosphorylation, but not total protein. P-values by paired t-test.

We next explored if cultured HGSC cells directly respond to IL-6 and/or IL-8 exposure. Using RT-qPCR to assay mRNA expression of IL-6 receptors (*IL6R* and *IL6ST*) and IL-8 receptors (*CXCR1* and *CXCR2*) in PEO1 cells (*TP53-/-*, *BRCA2*-mutant), we observed robust expression of *IL6ST*, but low or no expression of the other receptors **(Fig. 3G)**. In concordance with RT-qPCR, flow cytometry analysis showed that over 80% of PEO1 cells expressed surface IL6ST, but less than 1.5% expressed IL6R, CXCR1, or CXCR2 **(Fig. 3H** with representative flow gating shown in **Supplementary Fig. S6-7)**. We then incubated PEO1 cells directly in 50 ng/mL of recombinant IL-6, IL-8, or both for 15 minutes or 1 hour and performed immunoblot for resultant intracellular signaling, including phosphorylated STAT3 for IL-6 and phosphorylated ERK1/2 for IL-8. At 15 minutes, STAT3 phosphorylation at Y705 occurred in response to IL-6 alone and in combination with IL-8 **(Fig. 3I)**. At 1 hour, no p-STAT3 was observed **(Fig. 3J)**, indicating a transient response. IL-8 did not induce ERK1/2 phosphorylation, and we did not observe changes to total STAT3 or the β-actin loading control in PEO1 cells. To determine if PEO1 cells could respond to the range of IL-6 concentration observed in HSCDA conditioned media **(Fig. 3A-F)**, we incubated the cells for 15 minutes in a range of IL-6 doses from 5 pg/mL to 50 ng/mL **(Fig. 3K)**. By densitometry analysis of p-STAT3 bands, we observed progressively increased phosphorylation from 50 pg/mL upward. We observed comparable results in parallel experiments in two additional HGSC cell lines, OVCAR4 (*TP53-/-*, *BRCA1/2* wild-type) and OVCAR8 (*TP53-/-*, *BRCA1/2* wild-type*, ERBB2* mutant, *KRAS* mutant, *CTNNB1* mutant), including RNA and protein expression of receptors **(Supplementary Fig. S8A-C)**. We also observed upregulation of p-STAT3 in these lines in response to IL-6 at 15 minutes (and across multiple doses), but not at 1 hour **(Supplementary Fig. S8D-I)**. Overall, the primary response appeared to be from IL-6 driving a transient phosphorylation of STAT3.

We next expanded our interrogation of differential active signaling response following exposure to HSCDA and CMA media. We incubated HSCDA and CMA media from four independent human female donors (Donors 6-9) in triplicate on PEO1 cells for 24 h and then analyzed protein expression and phosphorylation by reverse-phase protein array (RPPA) **(Fig. 3L-R)**. The heatmap **(Fig. 3L)** shows proteins that were differentially regulated consistently across all four donors. These include HSCDA-induced upregulation of fibronectin (detailed in **Fig. 3M**), glyceraldehyde 3-phosphate dehydrogenase (GAPDH), and tissue transglutaminase 2 (TG2). We also observed increased serine phosphorylation of kinases AKT1, AKT2, and p21-activating kinase 4 (PAK4). Consistently downregulated proteins in HSCDA media, relative to cells in CMA media, included tumor suppressors such as the regulatory p85 subunit of PI3K (PI3K-p85), protein tyrosine phosphatase non-receptor type 12 (PTPN12), programmed cell death protein 4 (PDCD4), and glycogen synthase kinase 3 beta (GSK3β) (detailed in **Fig. 3N**). Notably, while phosphorylated forms of PAK4 **(Fig. 3O)** and AKT2 **(Fig. 3Q)** were found at higher levels, the total forms of these proteins **(Fig. 3P, R)** were not differentially expressed between treatment groups, highlighting specific differences in serine phosphorylation in the absence of overall protein changes. Overall, these data indicate that HSCDAs secrete a pro-tumorigenic milieu of factors, with the potential to alter protein signaling and expression in HGSC cells.

### Ablation of HSCDAs attenuates tumor burden in syngeneic murine models of HGSC

Human HSCDAs secrete greater amounts of inflammatory cytokines IL-6 and IL-8 than CMAs, and paracrine signaling in conditioned media alters protein expression and phosphorylation in HGSC cells **(Fig. 3)**. Both IL-6 and IL-8 increase the metastatic and invasive capabilities of HGSC cells [10, 20, 21]. Adipocytes also enhance tumor cell proliferation by providing energy through lipid transfer and stimulating HGSC cell fatty acid metabolism [9–11]. We hypothesized that ablation of HSCDAs in mice may reduce tumor progression in syngeneic HGSC models. To specifically ablate HSCDAs, irradiated recipient C57BL/6J mice were transplanted with hematopoietic stem cells (HSC) from donor mice in which a mature adipocyte-specific adiponectin gene promoter controls expression of an attenuated diphtheria toxin (DTA) gene [13]. This approach ablates any HSC that differentiates into a mature adipocyte by endogenous DTA expression, thereby specifically abrogating HSCDA production in the recipient mice. Importantly, CMAs are unaffected by myeloablation and transplant, as they are derived from host mesenchymal precursors that do not express DTA. We refer to this mouse cohort as “HSCDA Deficient” or HSCDA-Def. Control mice were generated by irradiation and transplant of HSCs from donor mice encoding a reporter gene but lacking DTA allowing the production of both HSCDAs and CMAs from hematopoietic and mesenchymal precursors, respectively. We refer to this mouse cohort as “HSCDA Proficient” or HSCDA-Pro. This is a complex procedure with potential confounders (e.g., x-ray irradiation and bone marrow transplant) that are uncommon in HGSC tumor studies. It cannot remove HSCDAs generated prior to irradiation, but no other methods currently exist to completely eliminate HSCDAs at this time. However, this procedure has been demonstrated to successfully reduce the HSCDA population from 16% down to <5% of adipose cells [13] and has been utilized to demonstrate the effects of even modest reduction in HSCDA abundance.

Following a six-week recovery from HSC transplantation, HSCDA-Def and HSCDA-Pro mice were engrafted with ID8 (*Tp53-/-*, *Brca2-/-*, GFP+Luc+) cells by intraperitoneal (IP) injection **(Fig. 4A)**. Prior studies have extensively characterized this syngeneic murine model as being representative of human disease, particularly in terms of its propensity to target and colonize the omentum before disseminating to other peritoneal sites [2, 22, 23]. Luciferase expression by ID8 cells was examined by IP luciferin injection and live imaging at 7, 21, and 35 days after cell injection. Total flux and percent change in flux (relative to 7 days post-ID8 injection) were similar between HSCDA-Pro and HSCDA-Def mice **(Fig. 4B-C)**. At 39 days post-ID8 injection, tumor-bearing mice were euthanized, and necropsy was performed to assess tumor burden and collect specimens for downstream analyses of the tumor and peritoneal environment. Representative necropsy images of the peritoneal space, including GFP+ omenta and disseminated tumors are shown in **Supplementary Fig. S9**. Average total omentum weight **(Fig. 4D)** strongly trended lower in HSCDA-Def mice compared to HSCDA-Pro mice (0.047 g vs 0.063 g; p=0.0921). The average number of disseminated tumor nodules **(Fig. 4E)** within the mesentery and other peritoneal locations was also reduced in HSCDA-Def mice (24.5 vs. 35.9; p=0.0566). The average total mass per mouse of these tumor nodules **(Fig. 4F)** was significantly lower in HSCDA-Def mice (0.089 g vs. 0.125 g; p=0.0484).

**Fig. 4.**
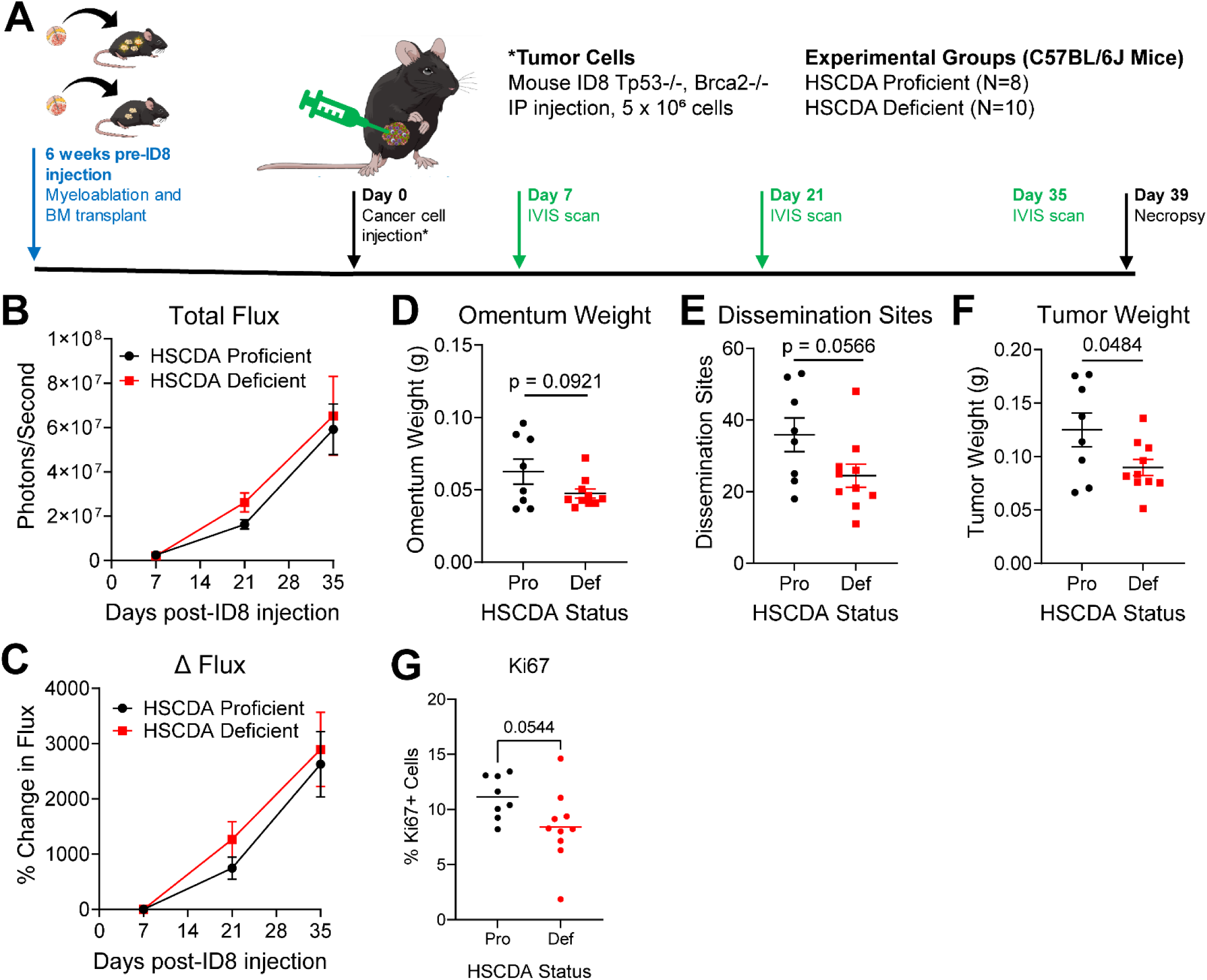
Ablation of HSCDAs reduces tumor burden in the ID8 syngeneic mouse model of HGSC. **(A)** Experimental timeline, including bone marrow transplant and subsequent tumor study. **(B)** Total flux and **(C)** percent change in total flux (relative to 7 days post-ID8 injection) by luciferase live imaging. **(D)** Weight of omental tumors. **(E)** Number of disseminated tumor nodules. **(F)** Weight of disseminated tumor nodules. For **(D-F)**, graphs show individual data points with mean ± SEM. p-values by unpaired t-test. **(G)** The percentage of Ki67-positive tumor cells was determined using QuPath software (representative images are shown in **Supplementary Fig. S10**). For **(G)**, individual data points with means (horizontal bar). p-values by unpaired t-test. Pro = HSCDA-Pro; Def = HSCDA-Def.

Given the reduced tumor burden in HSCDA-Def mice, we performed immunohistochemistry (IHC) on omental tumor sections to determine if tumor cell proliferation was affected by HSCDA status. Representative brightfield images of Ki67 (proliferation) are shown in **Supplementary Fig. S10**. Consistent with reduced tumor burden, the percentage of Ki67 positive tumor cells strongly trended lower (p=0.0544) in HSCDA-Def mice (average = 8.4%) than in HSCDA-Pro mice (average = 11.1%) **(Fig. 4G)**, suggesting that the HSCDA-depleted peritoneal adipose environment was less conducive to tumor growth. To examine the peritoneal cytokine milieu of tumor-bearing mice, we collected a portion of undiluted ascites fluid during necropsy. Cells were removed by centrifugation, and cytokine concentrations in cell-free ascites fluid were analyzed by multiplex ELISA: BDNF, IL-1β, IL-6, IL-10, Insulin, Leptin, MCP-1, and TNF-α. Ascites cytokine concentrations were not significantly different between mouse groups **(Supplementary Fig. S11)**, however we noted that IL-6 was higher in HSDCA-Def ascites (mean = 976.9 pg/mL) ascites than in HSCDA-Pro ascites (mean = 1214.9 pg/mL) (unpaired t-test p=0.0776).

As further confirmation of the effects of HSCDA ablation, we performed a follow-up tumor study in HSCDA-Pro and HSCDA-Def mice using the “SO” model [a kind gift from the Sandra Orsulic laboratory, UCLA]. SO cells are *Tp53-/-*, *Brca1/2* wild-type, *Hras^V12^* and *Myc* amplified. After an 8-week recovery from myeloablation and bone marrow transplant, SO cells were implanted by IP injection into recipient C57BL/6J mice. The SO model is extremely aggressive, so only 20,000 cells were injected per mouse. Accordingly, mice rapidly formed large solid omental and peritoneal tumors, necessitating the euthanization of all mice within 20 days post-SO injection. Consistent with the ID8 model, we observed non-significant reduction in omental tumor weight (p=0.29), as well as a strong trend toward reduced number (p=0.06) and mass (p=0.08) of non-omental disseminated peritoneal tumors in HSCDA-Def mice **(Supplementary Fig. S12)**. While the effects are modest in magnitude, the ID8 and SO studies collectively demonstrate that HSCDA ablation reduces overall intraperitoneal tumor burden in two independent syngeneic HGSC models.

### HSCDA ablation alters tumor transcriptional and immune microenvironment in ID8 tumor-bearing mice

We next assessed the HSCDA-dependent transcriptome and tissue architecture. We conducted RNA-Seq of ID8 omental tumors from HSCDA-Pro (n=4) and HSCDA-Def (n=4) mice to characterize the transcriptional profiles of the tumor microenvironment. As a measure of donor cell engraftment and expansion into recipient omentum, we assessed transcription of a tdTomato transgene, expressed specifically by donor HSCs in all recipient mice. We found that tdTomato expression, as a percentage of total transcriptional counts per mouse, was lower in HSCDA-Def mice **(Fig. 5A)**. We next examined differential gene expression. PCA showed broad clustering of HSCDA-Pro (blue) and HSCDA-Def (red) groups **(Fig. 5B)**. fGSEA analysis identified multiple differentially regulated Hallmark pathways **(Fig. 5C)**. Notably, three pathways we identified as enriched in human HSCDA cultures – Fatty Acid Metabolism, Oxidative Phosphorylation, and Adipogenesis **(Fig. 2F)** – were depleted in HSCDA-Def mice **(Fig. 5D-F)**. No tumor or ascites samples remained for direct measurement of metabolites; however, ORA pathway analysis of a human metabolomic and lipidomic dataset [24] showed significant differences in Fatty Acid Metabolism between human HGSC ascites fluid that contained low IL-6 (n=10) or high IL-6 (n=10) concentrations **(Supplementary Fig. S13)**.

**Fig. 5.**
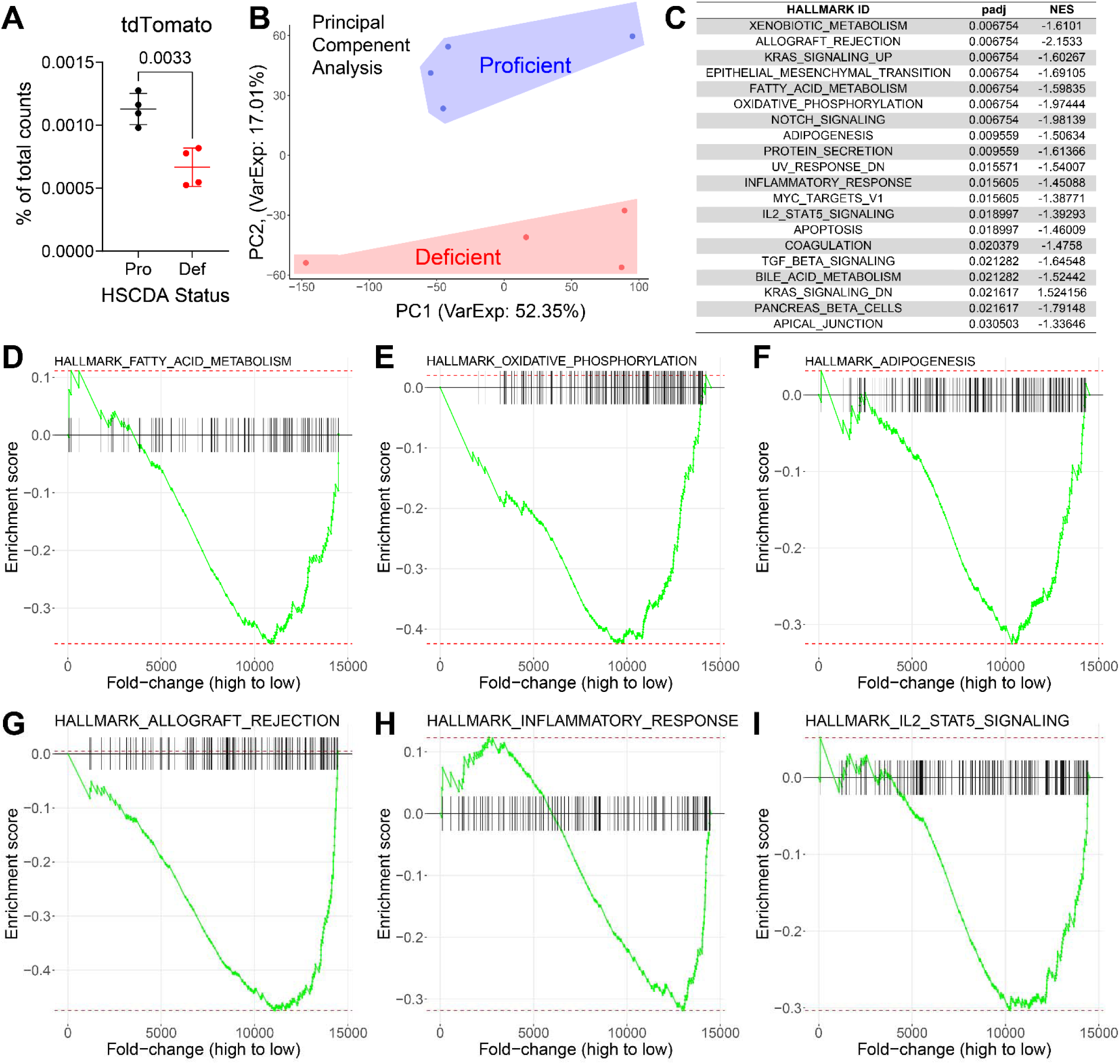
Ablation of HSCDAs in mice alters ID8 omental tumor gene transcription. **(A)** tdTomato (exogenous transgene from bone marrow donor cells) expression in ID8 omental tumor RNA-Seq from HSCDA-Pro (n=4) and HSCDA-Def (n=4) mice, expressed as percentage of total counts. Error bars = mean ± SD. Unpaired t-test. **(B)** Principal component analysis. **(C)** Significantly differentially regulated hallmark pathways by fGSEA. padj = p value adjusted for multiple comparisons. NES = Normalized Enrichment Score (adjusted for multiple comparisons). A positive NES indicates over-enrichment of pathway genes in HSCDA-Def mice relative to HSCDA-Pro mice, while a negative NES indicates under-enrichment in HSCDA-Def mice relative to HSCDA-Pro. **(D-F)** fGSEA enrichment plots for hallmark metabolic pathways. **(G-I)** fGSEA enrichment plots for hallmark immune pathways. The U-shaped plots in **D-I** correspond to negative NES in **C**.

fGSEA analysis also revealed the depletion of multiple immune-related Hallmark pathways, including Allograft Rejection, Inflammatory Response, and IL-2 STAT5 Signaling **(Fig. 5G-I)**. We then performed multispectral IHC (mIHC) on omental tumors to assess whether HSCDA status affected the tumor immune microenvironment. Tissue sections were stained using two separate panels of antibodies, both of which included staining for nuclei (DAPI) and the transcription factor WT1, which identifies ID8 tumor cells. The “innate” panel **(Table 2)** detected myeloid cells including dendritic cells (DC), natural killer cells (NK), macrophages, and subpopulations of myeloid derived suppressor cells (MDSCs). The “adaptive” panel **(Table 3)** detected lymphoid cells including B-cells and several subtypes of T-cells. After tissue and cell segmentation, cell densities were quantified in regions of interest. We found that the densities of DCs (CD11c+) and NKs (CD49b+/CD31-) were significantly lower in omental tumors from HSCDA-Def mice compared to HSCDA-Pro mice **(Fig. 6A-B)**. Proportions of other immune cell types (T-cells, MDSCs, and macrophages) in omentum tumors were unchanged by HSCDA status **(Supplementary Fig. S14)**. We also calculated the number of each immune cell type within 50 microns of WT1+ tumor cells. Within this radius, the numbers of DCs, NKs, and B-cells (B220+) were reduced in the omental tumors of HSCDA-Def mice compared to HSCDA-Pro mice **(Fig. 6C-E)**. Other cell types were unchanged **(Supplementary Fig. S14)**. Given the changes to DC and NK populations, we analyzed the activity status of DCs in a third mIHC panel **(Table 4)** using CD80+ and MHC II for mature dendritic cell activation, and Tbet (a transcription factor that activates IFN-γ) as a marker of active NK cells. The density of MHCII+ dendritic cells was reduced in tumors from HSCDA-Def mice **(Fig. 6G)**, while CD80+ DCs and Tbet+ NKs were unchanged **(Fig. 6F, H)**. These analyses show that HSCDA ablation affected not only tumor burden and bulk transcription but also the tumor immune microenvironment.

**Fig. 6.**
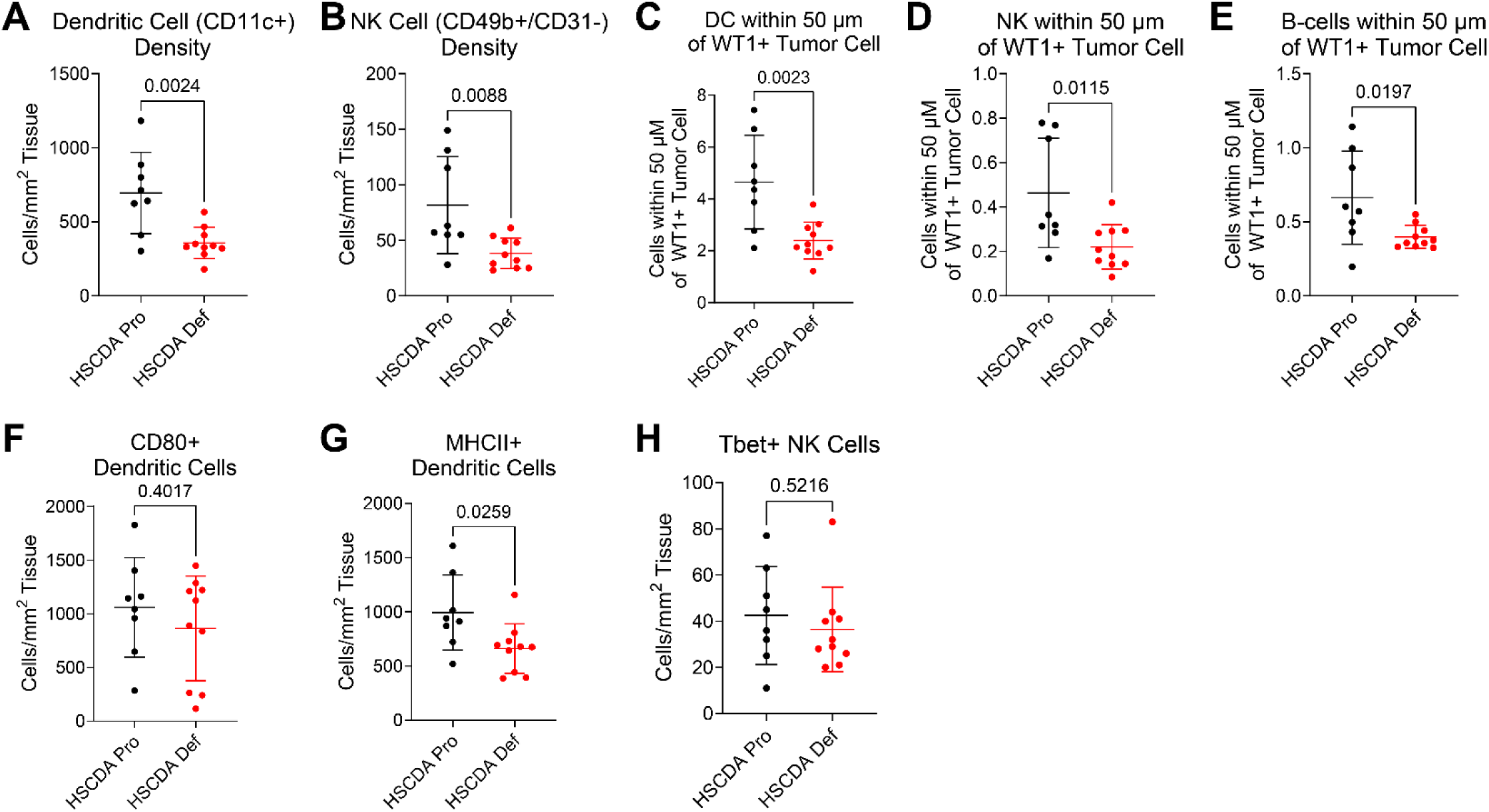
Ablation of HSCDAs reduces NK and DC density in ID8 omentum tumors. Omentum tumors from ID8-tumor bearing mice were analyzed by multispectral IHC for the indicated immune cells. **(A-B)** Cell densities of dendritic cells (CD11c+) and natural killer cells (CD49b+/CD31-) in omentum tumors. **(C-E)** Cell counts for the indicated DC, NK, or B-cells within 50 µm of a WT1+ tumor cell. **(F-H)** Cell densities of *mature* CD80+ or MHCII+ dendritic cells, or Tbet+ active NK cells. For all panels, bars indicate mean ± SD. Unpaired t-test.

**Table 2:** Multispectral IHC Antibody Panel for Innate/Myeloid Mouse Immune Cells.

| Antibody | Marker Expression |
| --- | --- |
| CD11b | Myeloid cells |
| CD11c | Dendritic cells |
| Ly6c | Monocytic MDSCs |
| Ly6g | Granulocytic MDSCs |
| CD49b | NK and endothelial cells |
| CD31 | Endothelial cells (negative on NK) |
| F4/80 | Macrophages |
| WT1 | ID8 tumor cells |
| DAPI | Nuclei |

**Table 3:** Multiplex IHC Antibody Panel for Adaptive/Lymphoid Mouse Immune Cells.

| Antibody | Marker Expression |
| --- | --- |
| CD3 | T-cells |
| CD4 | Helper T-cells |
| CD8 | Cytotoxic T-cells |
| Granzyme B | Effector T-cells |
| FOXP3 | Regulatory T-cells |
| B220 | B-cells |
| WT1 | ID8 tumor cells |
| DAPI | Nuclei |

**Table 4:** Multiplex IHC Antibody Panel for Mature/Active Dendritic and Natural Killer Cells.

| Antibody | Marker Expression |
| --- | --- |
| CD11c | Dendritic cells |
| CD80 | Co-stimulator (active DC) |
| MHC II | Antigen presentation (DC) |
| CD49b | NK and endothelial cells |
| CD31 | Endothelial cells (negative on NK) |
| Tbet | Transcription factor for IFN-γ (active NK cells) |
| DAPI | Nuclei |

## LIMITATIONS OF THE STUDY

Our *in vitro* HSCDA data are limited to the nine original adipose biopsy donors. It should be noted that these biopsies are from subcutaneous adipose depots, rather than visceral. While experiments were performed in three human cell lines that are widely accepted as high-quality models of HGSC [25], they cannot reproduce the entire gamut of potential genetic subtypes. We examined HSCDA and CMA conditioned media for cytokines, but other factors (e.g., metabolic factors, other signaling proteins, or exosomes) may also influence HGSC cell growth *in vitro* or tumor progression *in vivo*. Our *in vivo* data are derived from two high-quality syngeneic HGSC models but cannot encompass the variability of human disease. We have described transcriptional and immune changes in HSCDA-Def mice, which suggest potential pathways by which HSCDAs promote HGSC progression, however we do not yet have a direct cause-and-effect mechanism to explain reduced tumor growth. Ongoing and future experiments will generate omentum-derived HSCDAs and explore adipose-HGSC signaling pathways. These are currently beyond the scope of this initial characterization.

## DISCUSSION

Human data suggest that the proportion of HSCDAs increases over time [15]. In mouse models, HSCDAs preferentially accumulate in VAT and GAT depots, and their production is inversely correlated with estrogen signaling [17]. Crucially, most ovarian cancer diagnoses occur post-menopause [26], and the aged peritoneal microenvironment can be highly conducive to ovarian cancer progression [27]. We hypothesized that HSCDAs may promote HGSC progression by establishing a pro-tumor niche within VAT, such as the omentum, a site of high HGSC dissemination in human cases.

Human hematopoietic and mesenchymal adipocyte precursors are equally competent for adipogenic differentiation **(Fig. 1** and **Supplementary Fig. S2-S4)**; however, human HSCDAs have a distinct transcriptional profile compared to CMAs, characterized by the downregulation of Hallmark cell cycle pathways and upregulation of multiple metabolic pathways **(Fig. 2)**. ELISA of adipocyte conditioned media showed that human HSCDAs secrete higher amounts of IL-6 and IL-8 compared to CMAs **(Fig. 3A-F)**. Our RPPA data **(Fig. 3L-R)** indicated that HGSC PEO1 cells in HSCDA media, relative to CMA media, exhibit upregulated tissue transglutaminase 2, which induces epithelial-to-mesenchymal plasticity (EMP) and facilitates ovarian tumor dissemination [28, 29], as well as fibronectin, which is broadly associated with EMP in multiple cancer types [30]. Serine phosphorylation was also elevated in these cells, including pro-survival AKT1, AKT2, and PAK4, a group II serine/threonine protein kinase that can regulate cytoskeletal remodeling [31, 32] and Wnt/β-catenin signaling [33]. Interestingly, we observed that PEO1 cells cultured in HSCDA media exhibited downregulated GSK3β, a tumor suppressor and negative regulator of Wnt/β-catenin signaling. Other downregulated tumor suppressors included PTPN12, a tyrosine phosphatase that regulates PTEN and AKT signaling and suppresses the migration of ovarian cancer cells [34, 35], and PDCD4, a pro-apoptotic tumor suppressor with clinical significance in gliomas and colorectal cancers [36, 37].

While adipocyte-derived IL-6 and IL-8 have been shown to promote ovarian cancer survival, metastasis, and chemoresistance [10, 38], our data show that PEO1, OVCAR4, and OVCAR8 cells in culture express low levels of IL-6 receptor (IL6R) and IL-8 receptors (CXCR1 and CXCR2) **(Fig. 3** and **Supplementary Fig. S8)**. It is therefore likely that additional signaling mechanisms regulate the observed differences in HGSC cell proliferation and protein expression in HSCDA media. Prior studies have shown that exosomes from human omentum conditioned media can increase the proliferative and invasive capacity of ovarian cancer cells [39], and microRNAs from adipose tissue-derived exosomes can regulate gene expression in non-adipose tissues [40, 41]. Both PTPN12 and PDCD4, tumor suppressors that were downregulated in PEO1 cells in HSCDA media **(Fig. 3)**, are regulated by microRNA miR-503 [42, 43]. Future work examining the contribution of exosome-derived factors, including microRNAs, may reveal additional mechanisms for our observations.

Results from our *in vivo* studies in two independent syngeneic HGSC models showed that partial ablation of HSCDAs moderately reduced overall tumor burden, particularly by lowering the number and total mass of metastatic tumor nodules in the mesentery and other peritoneal sites **(Fig. 4** and **Supplementary Fig. S9** and **S12)**. Reduced Ki67 positivity in ID8 tumor cells from HSCDA-Def mice suggests that lower tumor burden may be due to reduced proliferation **(Fig. 4G** and **Supplementary Fig. S10A-B)**. Bulk RNA-Seq of ID8 omentum tumors revealed that HSCDA ablation downregulated metabolic pathways **(Fig. 5D-F)** that were conversely upregulated in human HSCDA cultures **(Fig. 2F-I)**, suggesting that HSCDA ablation contributed to a reduction in their expression in mice. Combined with the observed reduction in tumor burden and reduced Ki67, these data suggest that the peritoneal metabolic environment of HSCDA-Def mice was less supportive of tumor cell metastasis and growth **(Fig. 4G** and **Supplementary Fig. S10)**. No significant differences were observed in cytokine concentrations in cell-free ascites from ID8 tumor-bearing mice, although IL-6 was higher in HSCDA-Def mice **(Supplementary Fig. S11)**, which is contrary to expectations given the higher IL-6 concentrations secreted from human HSCDAs. Samples were insufficient for further lipid or metabolite analysis, but analysis of a human HGSC ascites metabolite dataset suggests that IL-6 can influence lipid metabolic pathways **(Supplementary Fig. S13)**. Further tumor studies are planned and underway to directly characterize the visceral metabolic environment of HSCDA-Pro and HSCDA-Def mice, including free fatty acids and oxidation products in tumor tissue and ascites fluid.

RNA-Seq of omentum ID8 tumors also revealed downregulation of multiple immune-related pathways **(Fig. 5G-I)** in HSCDA-Def mice relative to HSCDA-Pro mice. Omental tumors in HSCDA-Def mice showed reduced overall density of DCs and NKs, as well as fewer DCs, NKs, and B-cells within a 50 µm radius of tumor cells **(Fig. 6A-E)**. CD80+ DC density was unchanged between mouse groups, indicating similar DC maturation **(Fig. 6F)**. However, MHCII+ DC density was reduced in HSCDA-Def mice, suggesting differences in antigen presentation **(Fig. 6G)**. IL-6 and IL-10 can suppress MHC II presentation on DCs [44, 45]. IL-6 and IL-10 concentrations were not significantly different between mouse groups in cell-free ascites, but the mean concentration of IL-6 in HSCDA-Def mice did trend higher **(Supplementary Fig. S11)**. The reduced tumor burden and proliferation in HSCDA-Def mice may have also provided less tumor antigen for presentation. Tbet+ NK density was unchanged between mouse groups, suggesting similar cytotoxic activity **(Fig. 6H)**. A caveat of these findings is the difference in tdTomato expression between HSCDA-Pro and HSCDA-Def omentum tumor samples **(Fig. 5A)**, which may reflect a difference in engraftment of hematopoietic cells. However, significant differential engraftment would likely result in an overall difference in all myeloid populations, which was not observed. In terms of function, innate immune cells such as NKs and DCs can directly target malignant cells and broadly promote the recruitment of adaptive immune cells, and it has been proposed that B-cells also promote the HGSC anti-tumor response [46]. However, tumor-promoting roles have also been described for each of these cells. For example, tumor microenvironment components can suppress NK cell cytotoxicity and induce angiogenesis [47]. In a study of murine adenocarcinoma and melanoma, NK cells rapidly lost effector functions after entering the tumor environment [48]. Additionally, regulatory B-cells can suppress immune response through the production of anti-inflammatory cytokines [49, 50]. Further exploration is needed to characterize the populations of tumor infiltrating DCs, NKs, and B-cells, and whether the balance of pro- and anti-tumor cells can be shifted by the presence of HSCDAs.

In conclusion, we present the first study characterizing human HSCDAs in the context of ovarian cancer, as well as two independent *in vivo* tumor studies comparing HSCDA-Def and HSCDA-Pro mice. Our results suggest HSCDAs are important contributors to ovarian cancer progression that may alter the peritoneal immune and metabolic environment to establish a metastatic niche and promote disease progression. Future studies are underway to shed light on specific mechanisms. Use of genetically engineered murine models that develop spontaneous cancers [51, 52] may prove useful, as they give a longer period for the pro-tumorigenic properties of HSCDAs to become apparent. The current models, while showing trends and some significant differences, are limited to an acute tumor-adipose exposure time of only a few weeks. Despite this limitation, our findings in mice raise the possibility of specifically targeting HSCDAs in humans as a method of reducing metastatic tumor burden.

## RESOURCE AVAILABILITY

Requests for further information and resources should be directed to and will be fulfilled by the lead contact, Zachary L Watson.

- RNA-Seq data are available at GEO at the accession numbers listed in the materials and methods.
- Other raw data are available at Figshare: https://doi.org/10.6084/m9.figshare.34112514 or Mendeley: 10.17632/spjzn5wxzk.1
- Human HGSC cell lines are listed by RRID and are available from ATCC or other repositories.
- Syngeneic mouse lines are available from their originating laboratories (ID8: McNeish lab [53]; SO: Orsulic Lab, UCLA).
- Human adipose precursors are restricted to research use at the University of Colorado by protocols of the Colorado Multiple Institutional Review Board (COMIRB).
- Mouse strains are listed by RRID and are available from Jackson Labs.
- Primer sequences generated in this study are listed in the supplementary materials and may be purchased commercially.
- All other reagents such as media, growth factors, chemicals, cytokines, and antibodies are listed and are commercially available.
- No proprietary code was generated by this study. All tools for RNA-Seq analysis are listed and are available for public use.

## MATERIALS AND METHODS

### HGSC cell culture

PEO1 (RRID: CVCL_2686), OVCAR4 (RRID: CVCL_1627), and OVCAR8 (RRID: CVCL_1629) were obtained from the Gynecologic Tissue and Fluid Bank (GTFB) at the University of Colorado and were authenticated at the University of Arizona Genomics Core using short tandem repeat DNA profiling. Cells were repeatedly confirmed free of mycoplasma using MycoLookOut PCR (Sigma). Standard culture condition for these lines is 37 °C in 5% CO_2_ in RPMI 1640 supplemented with 10% fetal bovine serum (FBS) and 1% penicillin/streptomycin.

### Study Participants (reproduced with author permission from [18])

The study was approved by the Colorado Multiple Institutional Review Board (COMIRB) and written informed consent was obtained from all participants. All procedures took place at the Clinical and Translational Research Center (CTRC) at the University of Colorado Anschutz Medical Campus (CU-AMC). Samples were obtained from participants enrolled in NCT02654925, NCT04043520, and NCT03396978. Participants were healthy males and females either 20-50 or >50-75 years of age (y). Older women were postmenopausal as identified by no menstrual cycles within the last 12 months and confirmed by follicle-stimulating hormone (FSH)≥ 30 mIU/mL. Premenopausal women were not currently pregnant or lactating. Inclusion criteria included a BMI between 20 and 40 kg/m^2^. Exclusion criteria included current (within the last 6 months) use of any hormone replacement or hormonal contraceptive, use of glucose lowering medication and diagnosis of type 2 diabetes or uncontrolled metabolic disorders.

### Adipose tissue biopsy collection and processing (reproduced with author permission from [18])

The skin and subcutaneous abdominal adipose tissue directly adjacent to the umbilicus underwent local numbing with 1% lidocaine without epinephrine, after which the adipose tissue samples were obtained using a modified Coleman’s manual vacuum (“mini liposuction”) technique. A small incision was made, followed by infiltration of the adipose tissue with ∼30- 50 mL of 0.15% tumescent lidocaine solution in 0.9% normal saline using a Coleman infiltration cannula. Approximately one to three grams of adipose tissue was removed using a Coleman aspiration cannula. Adipose tissue was rinsed with collection buffer (Krebs-Ringers-HEPES (KRH), 2.5 mM glucose, 200 nM adenosine, pH 7.4) and obvious blood vessels and clots removed. 250-500 mg was flash frozen in liquid nitrogen and stored at -80 °C. Remaining tissue was transported to the laboratory where collection buffer was replaced with prewarmed digestion buffer (KRH, 2.5 mM glucose, 2% fetal bovine serum [FBS], 200 nM adenosine, 1mg collagenase/0.25 g tissue, pH 7.4). Tissue was digested for 30-60 minutes in a shaking water bath maintained at 37 °C. The resultant cell suspension was filtered through 250 µm mesh and digestion stopped with 1X volume of wash buffer (Hanks Balanced Salt Solution [HBSS] + 2% FBS + 200 nM adenosine). Samples were spun at 300 x *g* for 10 min, the floating adipocytes removed to a clean tube, and wash steps repeated. The stromal pellets were combined and spun at 500 x *g* for 10 min to pellet the cells. Here forward, all spins were at 500 x *g* for 10 min unless otherwise stated. The stromal cell pellet was resuspended in 10 mL Erythrocyte Lysis Buffer and incubated for 10 min on ice. Simultaneously, 1 mL of blood underwent red blood cell lysis. When applicable, adipose and blood samples were processed in parallel from here forward. After erythrocyte lysis, samples were spun and cell pellet resuspended in 1 mL wash buffer. Live cells were counted via trypan blue detection on a Cellometer X1 (Nexcelom Bioscience LLC, Lawrence, MA) cell counter.

### Fluorescence activated cell sorting (FACS) of processed adipose biopsies (reproduced with author permission from [18])

After counting, cells were spun and resuspended at a concentration of 1 × 10^6^ cells/100 µL in wash buffer. Human TruStain FcX was added at 5µL/100µL cell suspension and cells incubated on ice for 10 min. All lineage antibodies were added at 5µL per million cells in 100 µL staining volume (or 0.25µg/10^6^ cells) and mixed gently. Samples were incubated on ice in the dark for 25 min. Following incubation, samples were spun, the supernatant aspirated, and the cell pellet resuspended in wash buffer. After removal of the second supernatant, the cells were resuspended in flow buffer (HBSS + 5% FBS + 200 nM adenosine, pH 7.4) with volumes adjusted to 500µL – 1mL for cell sorting. DAPI was added to a final concentration of 1 mg/mL. Cell sorting was initiated within 15 minutes. Unstained cells, single color compensation beads, and fluorescent minus one (FMO) controls were used. Antibodies are listed in the supplementary information of reference [18]. Stromal vascular fraction cells were sorted using an Astrios cell sorter with Summit 6.3 software (Beckman Coulter, Fullerton, CA, USA). A 70 µm nozzle tip was used with a sheath pressure of 60 PSI and a drop drive frequency of approximately 95,000 Hz with an amplitude of 15V. Manufacturer’s protocols were followed to quality control and set up the sorter. The sheath fluid was IsoFlow. The sample and collection tubes were maintained at 10 °C using an attached recirculating water bath. To keep cells in suspension, the Astrios’ SmartSampler sample station was set to maintain an agitation cycle of 5 seconds on and 8 seconds off. The sample flow rate was set to a pressure differential of <1.0 PSI. Sort mode was set to Purify 1. Appropriate signal compensation was set using single-color control samples. Cells were identified with the following surface markers: mesenchymal lineage cells, CD45NEG; conventional mesenchymal adipocyte precursor cells, CD45NEG/CD31NEG/CD29POS; hematopoietic Lineage, CD45POS; myeloid lineage, CD14POS; macrophages (ATM) & monocytes, CD14POS/HLA-DRPOS with divergent expression of CD206 (ATM CD206POS, monocytes CD206NEG); neutrophils, CD14POS, CD16POS, HLA-DRNEG, Siglec-8NEG; basophils, CD14NEG/Siglec-8POS; lymphocytes, CD14NEG/CD206NEG.

### Expansion of sorted cells (reproduced with author permission from [18])

Sorted cells were gently vortexed and spun for 10 min at 500 x *g*. Cell pellets were resuspended in growth medium and conventional mesenchymal lineage cells were plated directly onto cell culture treated plastic plates for proliferation by standard techniques at a density of ∼4,500 cells/cm2. Fresh growth media was applied the next day, and media changes took place every 2-3 days throughout the studies. Cells were grown to 70-80% confluence, detached with 30-60 min room temperature incubation in Accutase and passaged 1-2 additional times. Cells were frozen down in liquid nitrogen for use in future experiments. ATM/monocytes, neutrophils, and lymphocytes were each resuspended in 10 mL growth media at 40,000 cells/µL. Fibrinogen (60 µL, 5 mg/mL in 0.9% saline) was added, mixed gently and transferred to one well of a 96 well plate (per 400,000 cells) containing 1.4 µL of thrombin (50 U/mL) and the suspension mixed by gentle pipetting. Clots formed over 30–60 minutes and growth media was subsequently overlaid. Media was refreshed every 2-3 days throughout the study. Cells were recovered from the clots after 5-7 days. The culture medium was removed and matrices rimmed with a 27-gauge needle. The remaining liquid in the well was removed and 100 µL of digestion mix (75 µL growth medium + 23 µL bovine plasminogen (1.12 mg/mL) + 2µL urokinase (20 U/µL)) was added for 60 minutes at 37 °C, mixing by pipetting every 20 minutes. After digestion, cells were pelleted in a 1.5 mL tube at 500 x *g* for 10 minutes, resuspended in growth medium, and re-plated directly onto plastic into the same number of wells of a 96 well plate. All cells then underwent identical culture methodology to the mesenchymal lineage cells.

### Primary preadipocyte culture, adipogenic differentiation, and conditioned media collection

Primary mesenchymal and myeloid-derived adipocyte precursors were grown in 15 cm tissue culture plates in Growth Media (GM): Alpha-MEM (Corning #10-022-CV) supplemented with 10% FBS (Gemini #100-106) and 0.1% penicillin/streptomycin (Sigma #P4333). Upon confluence, cells were differentiated for seven days in Complete Differentiation Media (CDM): DMEM/F12 (Corning #10-092-CV) supplemented with 33 μM D-biotin (Sigma #B4639), 17 μM pantothenate (Sigma #P5155), 1 μM rosiglitazone (Fisher #NC9331406), 0.5 mM 3-isobutyl-1-methylxanthine (IBMX; Sigma #I5879), 2 nM 3,3’,5-triiodo-L-thyronine (T3; Sigma #T6397), 10 μg/mL transferrin (Sigma #T8158), and 100 nM insulin (Sigma #I9278). Cultures were then maintained for seven days in adipocyte maintenance media (AMM): DMEM/F12 supplemented with 33 μM D-biotin, 17 μM pantothenate, and 10 nM insulin (same suppliers as CDM). At the end of the seven-day AMM period, media was removed from adipocyte cultures. Cells were gently washed with PBS, and then media was replaced with fresh AMM. After a 24-hour incubation, conditioned media was collected for subsequent use.

### Bulk RNA-Seq of human primary HSCDAs and CMAs

Total RNA was isolated from human primary HSCDA and CMA cultures using the RNeasy Plus kit (Qiagen). Library preparation and sequencing were performed by the University of Colorado Cancer Center Genomics Shared Resource (RRID:SCR_021984). Reads were analyzed by the University of Colorado Cancer Center Bioinformatics Core Facility (RRID:SCR_021983). RNA-Seq data were processed using the nf-core RNAseq pipeline (version 3.12.0; [54]). Briefly, Illumina adapters were removed using Cutadapt (version 3.4; [55]) as part of the trimgalore (0.6.10) package [56]. Reads were aligned using STAR (version 2.7.9a; [57]) to the human transcriptome (GRCh38, gene annotation from Ensembl release 104) and quantified using Salmon (version 1.10.1; [58]). Raw data with counts by gene was generated using tximport ([59]) on salmon quantified data. Normalized data was generated to counts per million [60]. Differential expression was calculated between sample groups using the limma R package [61]. Gene set enrichment analysis (GSEA) was performed using fold-change and the fgsea R package [62] with Hallmark gene sets from the Molecular Signatures Database [63], which were downloaded using the msigdbr R package. Raw and processed RNA-Seq data was deposited in the Gene Expression Omnibus (GSE277309).

### Multiplex ELISA

For human cytokines in conditioned media, AMM was incubated on HSCDA or CMA cells for 24 h, then collected. The concentration of 10 cytokines (IFN-γ, IL-1β, IL-2, IL-4, IL-6, IL-8, IL-10, IL-12p70, IL-13, and TNF-α) in conditioned media was measured using the V-PLEX Proinflammatory Panel 1 Human Kit (Meso Scale Discovery #K15049D). For mouse adipokines, ascites fluid was collected from the peritoneal space of tumor-bearing HSCDA-Pro and HSCDA-Def mice using a needle and syringe. The supernatant was cleared of cells by centrifugation, and the concentrations of 8 adipokines (BDNF, IL-1β, IL-6, IL-10, Insulin, Leptin, MCP-1, and TNF-α) were measured in the supernatant using the U-PLEX Adipokine Combo 1 mouse kit (Meso Scale Discovery #K15299K).

### Reverse-transcriptase quantitative PCR (RT-qPCR)

RNA was isolated from cells using the RNeasy Plus kit (Qiagen). mRNA expression was determined using SYBR green Luna One Step RT-qPCR Kit (New England BioLabs) on a Bio-Rad C1000 Touch thermocycler. Expression was quantified by the ΔCt method using target-specific primers and glyceraldehyde 3-phosphate dehydrogenase (*GAPDH*) control primers. mRNA-specific primers were designed to span exon-exon junctions to avoid the detection of genomic DNA. Primer sequences are shown in **Supplementary Table ST1**.

### Flow cytometry for ovarian cancer cell cytokine receptor surface expression

One million HGSC cells grown in standard conditions (see above) were collected and resuspended in 100 µL of flow assisted cell sorting buffer (FACS buffer: Hank’s Balanced Salt Solution [Corning #21-022-CV] with 2% FBS) and incubated in a 1:100 dilution of primary antibody for 20 minutes at 4 °C. Cells were then washed in 300 µL FACS buffer and centrifuged at 300 x *g* for 5 minutes. Cells were resuspended 300 µL FACS buffer and transferred to 5 mL tubes through 35 µm strainer caps (Falcon #352235). Full details of antibodies and isotype controls are given in **Supplementary Table ST2**. An Agilent Novocyte Penteon was used for detection, and data were analyzed using FlowJo 10.8.1.

### Immunoblotting

HGSC cells were seeded to sub-confluence in 10 cm dishes in standard media and allowed to settle overnight. Media was aspirated and cells were washed with PBS. Media was then replaced and supplemented with vehicle control, recombinant human IL-6 (BioLegend #570804), recombinant human IL-8 (BioLegend #574204), or both. Cells were incubated for 15 minutes or 1 hour, then rinsed and collected. Cell pellets were lysed in RIPA buffer (150 mM NaCl, 1% TritonX-100, 0.5% sodium deoxycholate, 0.1% SDS, 50 mM Tris pH 8.0) supplemented with protease inhibitor (Roche #11873580001) and phosphatase inhibitor (Roche #04906845001). Protein was separated by SDS-PAGE and transferred to PVDF membrane using the TransBlot Turbo (BioRad). Membranes were blocked in LI-COR Odyssey buffer (#927-50000) for use with LI-COR secondary antibody or in 2% BSA in TBS-T for use with HRP-linked secondary antibody for 1 hour at room temperature. Primary antibody incubation was performed overnight in blocking buffer at 4 °C. Membranes were washed 3 times for 10 minutes each in TBST (50 mM Tris pH 7.5, 150 mM NaCl, 0.1% Tween-20), then secondary antibodies were applied in blocking buffer for one hour at room temperature. Membranes were washed again 3 times for 10 minutes each in TBST. For LI-COR secondary antibody, bands were visualized using the LI-COR Odyssey Imaging System. For HRP-linked secondary antibody, membranes were incubated for 5 minutes in SuperSignal West Femto Maximum Sensitivity Substrate (Thermo #34095) and bands were visualized using a Syngene G:BOX. Densitometry of p-STAT3 bands was performed in ImageJ v1.54. Full details of primary and secondary antibodies are given in **Supplementary Tables ST3** and **ST4**.

### Generation of HSCDA-Pro and HSCDA-Def Mice

The University of Colorado Institutional Animal Care and Use Committee (IACUC) approved all mouse studies. Tumor burden analysis was performed in bone marrow transplanted (recipient) wild-type C57BL/6J mice (Jackson Labs, strain #000664; RRID: IMSR_JAX:000664). HSCDA-Def mice were generated by transplanting recipient mice with bone marrow from transgenic donor mice in which an adipocyte specific adiponectin gene promoter (AdipoQ-Cre) controls the expression of an attenuated diphtheria toxin (DTA). HSCDA-Pro mice were transplanted with bone marrow from donor mice encoding AdipoQ-Cre but lacking DTA. Prior to transplant, 8-to 12-week-old female recipient mice were myeloablated by a split 12 Gy dose of X-ray irradiation (6 Gy doses split by 4 hours). Irradiated mice were rescued with retroorbital injection of 1 × 10^6^ donor cells to reestablish the hematopoietic niche. Donor cells were obtained from fresh bone marrow of transgenic animals which can produce mature adipocytes (AdipoQ-Cre) or lack mature adipocytes (AdipoQ-Cre-DTA). Adipocyte-deficient donor mice were generated by crossing animals in which Cre recombinase expression is directed by the adiponectin promoter (Jackson Labs #010803, B6;FVB-Tg(AdipoQ-cre)1Evdr/J; RRID: IMSR_JAX:010803) with mice in which the DTA gene is placed downstream of a floxed stop codon in the ROSA26 locus (Jackson Labs #010527, B6;129-Gt(ROSA)26Sortm1(DTA)Mrc/J; RRID: IMSR_JAX:010527). Cre-mediated excision of the stop codon initiates DTA expression and cell death, which is restricted to mature adipocytes by the adiponectin promoter. Adipocyte proficient donor mice are the result of the same cross, excluding the DTA gene. Recipient mice recovered from bone marrow transplantation 6-8 weeks prior to tumor inoculation and experimentation protocols.

### ID8 Animal Study

Six weeks after bone marrow transplant, 5 million ID8 cells (*Tp53-/-*, *Brca2-/-*, GFP-luciferase+,[53, 64]) were implanted into HSCDA-Pro and HSCDA-Def mice by intraperitoneal injection. Tumor development was monitored in all mice via biweekly In Vivo Imaging System (IVIS) scanning, as described previously [65]. 39-days post-ID8 cell injection, mice were euthanized, ascites fluid was collected, and necropsy was performed to assess tumor burden. All tumors were collected and weighed, including omentum tumors and disseminated tumor nodules within the mesentery and other peritoneal tissues.

### SO Animal Study

Eight weeks after bone marrow transplant, 20,000 SO cells (*Tp53-/-*, *Brca1/2* wild-type, *Hras^V12^* and *Myc* transformed) were implanted into HSCDA-Pro and HSCDA-Def mice by IP injection. Mice were euthanized 20-days post-SO cell injection and necropsy was performed to assess tumor burden. All tumors were collected and weighed, including omentum tumors and disseminated tumor nodules within the mesentery and other peritoneal tissues.

### Bulk RNA-Seq of ID8 Murine Omentum Tumors

Library preparation and sequencing were performed by the University of Colorado Cancer Center Genomics Shared Resource (RRID:SCR_021984). Reads were analyzed by the University of Colorado Cancer Center Bioinformatics Core Facility (RRID:SCR_021983). RNA-Seq data were processed using the nf-core RNAseq pipeline (version 3.14.0; [66]). Briefly, Illumina adapters were removed using Cutadapt (version 3.4; [55]) as part of the trimgalore (0.6.7) package [56]. Reads were aligned using STAR (version 2.7.9a; [57]) to the Ensembl mouse transcriptome (GRCm39 release 104) and quantified using Salmon (version 1.10.1; [58]). Raw data with counts by gene was generated using tximport [59] on salmon quantified data. Normalized data was generated to counts per million [60]. Differential expression was calculated using the limma R package [61] comparing HSCDA-Def to the HSCDA-Pro sample groups. Volcano plot was generated with GraphPad Prism (ver. 10.2.3). Raw and processed RNA-Seq data was deposited in the Gene Expression Omnibus (GSE268528).

### Multiplex Immunohistochemistry (mIHC) and Image Analysis

Tissues were formalin fixed for 48 hours at room temperature, embedded in paraffin blocks using a tissue processor, and sectioned at 4 microns onto charged glass slides. Slides were deparaffinized, heat treated in antigen retrieval buffer, blocked, and incubated with primary antibodies (**Tables 2-4**), followed by horseradish peroxidase (HRP)-conjugated secondary antibody polymer, and HRP-reactive OPAL fluorescent reagents. The slides were stripped between each consecutive staining cycle with heat treatment in antigen retrieval buffer. Whole slide scans were collected with PhenoImager HT v2.0.0 software using the 20x objective and a 0.5-micron resolution. Regions of interest (ROIs) were selected and re-scanned using the 20x objective and the multispectral imaging cube. Spectral references and unstained control images were measured and inForm software v3.0 was used to create a multispectral library reference. The multispectral images (.im3 files) were spectrally unmixed and analyzed with tissue segmentation, cell segmentation, and phenotyping using inForm software v3.0 (Akoya Biosciences). Exported data were compiled and summarized using PhenoptrReports (Akoya Biosciences). Full antibody details are given in **Supplementary Table ST5**.

### Brightfield IHC and Image Analysis

Tissues were formalin fixed for 48 hours at room temperature, then paraffin embedded in blocks using a tissue processor and sectioned onto glass slides. All slides were de-identified and whole slide scans were collected on Akoya Biosciences Vectra 3 scanner using the 20x objective by the Human Immune Monitoring Shared Resource (RRID:SCR_021985). The slides were then analyzed on QuPath (v. 0.5.1 x64). ROIs were drawn to encompass the whole tissue, and background correction was done for H-DAB. For each stain, 4 ROIs were drawn for training sets and excluded from analysis for which cell and positive cell detect parameters could be identified. Cell detections were based on hematoxylin signal, with exclusions for minimum cell area and the inclusion of a median filter radius. Cell positivity was determined on DAB pixel intensity, and the percentage of positive cells was reported as the ratio of cells above a set DAB optical density mean to the total number of cells in the tissue. Data were exported and analyzed in GraphPad Prism 10 v. 10.3.1. The analyst was blinded to treatment group information until after the analysis was complete to avoid bias.

## Supporting information

All Supplemental Figures and Tables

## Ethics approval and consent to participate

All mouse studies were approved by the University of Colorado Institutional Animal Care and Use Committee (IACUC; protocols 157 [generation of HSCDA-Pro and HSCDA-Def mice] and 569 [syngeneic models of ovarian cancer]). The samples utilized for primary preadipocyte culture were obtained under approval by the Colorado Multiple Institutional Review Board (COMIRB) with written informed consent from all participants. Samples were obtained from participants enrolled in NCT02654925, NCT04043520, and NCT03396978 and shared in a de-identified manner as secondary use in this investigation as approved by COMIRB #22-0809.

## Consent for publication

N/A

## Conflict of interests

The authors declare that they have no conflict of interest.

## Funding

ZLW is a Scholar of the Colorado Specialized Center for Research Excellence (CO-SCORE) on Sex Differences (U54 AG062319) focused on Bioenergetic and Cardiometabolic Consequences of the Loss of Gonadal Function. This work was primarily supported by the Department of Defense Ovarian Cancer Research Program (ZLW, OC210257; SO, HT94252410193) and the Colorado Clinical & Translational Sciences Institute (ZLW, CCTSI CO-Pilot CO-J-20-006), with additional support by the merged Ovarian Cancer Research Alliance / Rivkin Center for Ovarian Cancer Research (ZLW, Pilot Award RPG-R-2024-2-1269446) as well as Institutional Research Grant #22-154-59 from the American Cancer Society to the University of Colorado Cancer Center (ZLW). Further support was provided by the American Cancer Society (BGB, RSG-19-129-01-DDC), the NCI (BGB, R37CA261987; ZLW, R03CA249571), NIDDK (KMG, K01DK109053), NIA (KMG and DJK, U54 AG062319), the VA (SO, I01BX006020), and the Boettcher Foundation. Use of the University of Colorado Bioinformatics and Biostatistics Shared Resource (BBSR), Human Immune Monitoring Shared Resource (HIMSR), and the Cell Technologies Shared Resource (CTSR) was supported by NCI grant P30CA046934 and by NIH/NCATS Colorado CTSA grant UL1TR002535. This work was supported by the Alpine HPC system, which is jointly funded by the University of Colorado Boulder, the University of Colorado Anschutz, Colorado State University, and the National Science Foundation (award 2201538).

## Author’s contributions

ERW (co-first author): performed experiments, analyzed data, prepared manuscript

CAB (co-first author): performed experiments, analyzed data, prepared manuscript FT

(co-first author): performed experiments, analyzed data, revised manuscript

VM: performed experiments, analyzed data

JKM: performed experiments

TMS: performed experiments

MPB: analyzed data, revised manuscript

MSR: provided study materials

RV: provided study materials, aided in study design

BC: provided study materials, revised manuscript

KRJ: analyzed data

KMG: provided study materials, aided in study design, revised manuscript

DJK: provided study materials, aided in study design

SO: provided study materials

BGB: provided study materials, aided in study design, data analysis and interpretation, revised manuscript

ZLW: designed the study, performed experiments, analyzed and interpreted data, prepared manuscript

## Additional Acknowledgments

The authors acknowledge with extensive gratitude the anonymous donors who provided biopsies for use in generating adipocyte cultures. We acknowledge philanthropic contributions from the Kay L. Dunton Endowed Memorial Professorship in Ovarian Cancer Research, the McClintock-Addlesperger Family, Karen M. Jennison, Don and Arlene Mohler Johnson Family, Michael Intagliata, Duane and Denise Suess, Mary Normandin, and Donald Engelstad. The corresponding author (Zachary L Watson) wishes to dedicate this manuscript in memory of Maggie Watson (1981-2022) and Mira Watson (2016-2022).

## SUPPLEMENTARY MATERIALS

Supplementary Figures S1 – S14

Supplementary Tables ST1 – ST5

The supplementary materials are prepared as a separate file as required by the instructions for authors.

## Notes

### Competing Interest Statement

The authors have declared no competing interest.

### Summary of Updates

Updated and simplified PCR analysis in Fig. 1. Now performed in triplicate and added to new Supplementary Fig. S5. Added further description of the mouse HSCDA ablation procedure, as well as limitations, caveats, and confounders. New analysis of a human HGSC ascites dataset (Supplementary Fig. S13). A new multispectral IHC panel to assess maturity and antigen presentation in DCs and maturity/cytotoxicity of NKs (new Fig. 6). Due to new data, all multispectral IHC data were moved to new Fig. 6. Table 4 was added for the new DC/NK panel. Supplementary Figures and Tables were added with new data, or were moved. The text was extensively edited to ensure that all figure and table references are correct for this new version.

