## Supplementary material for "Ablation of hematopoietic stem cell derived adipocytes reduces tumor burden in syngeneic mouse models of high-grade serous carcinoma": All Supplemental Figures and Tables

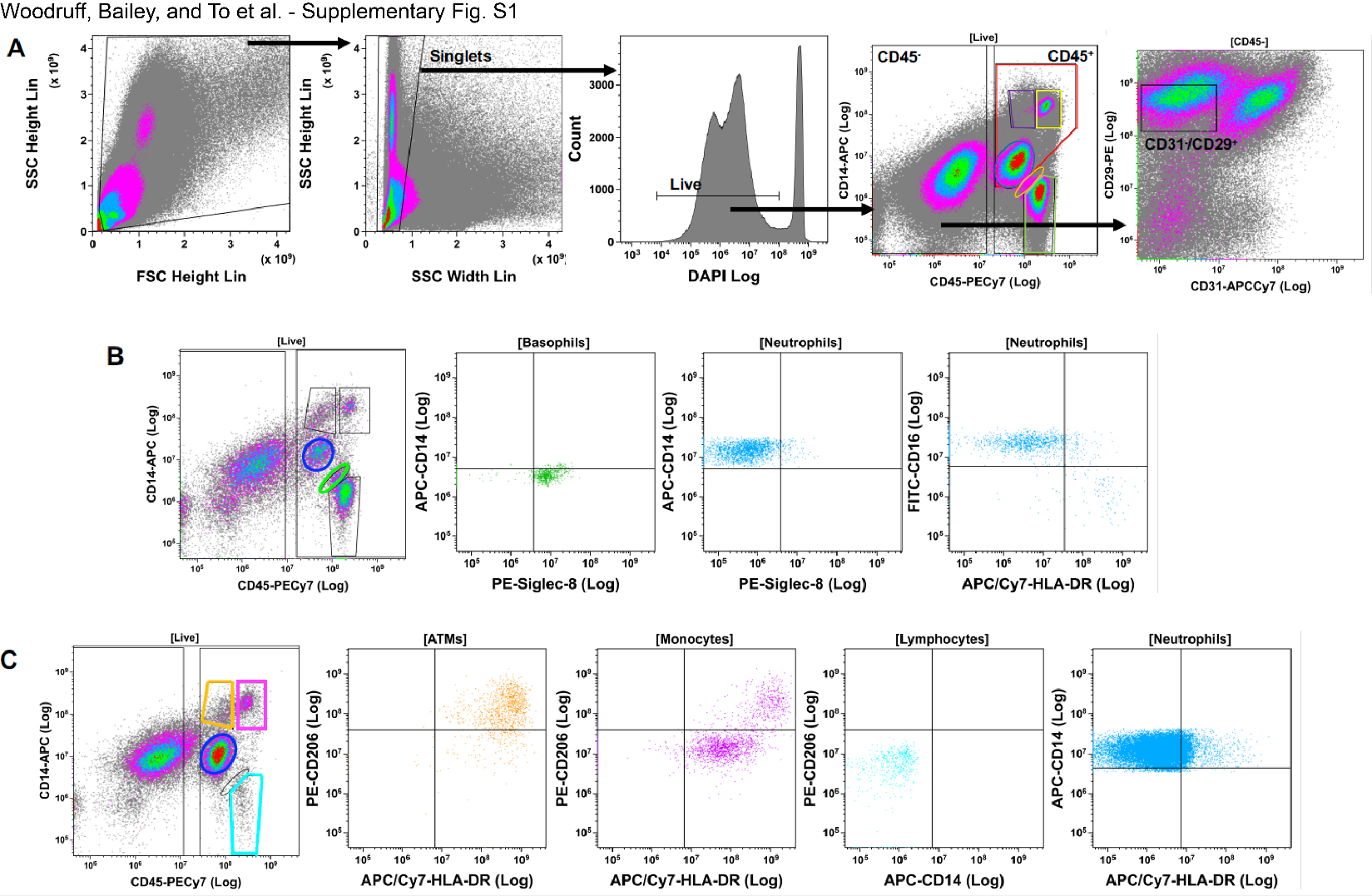

**Fig. S1. Flow Cytometry/Fluorescent Activated Cell Sorting (FACS) strategies for identifying mesenchymal and hematopoietic lineage cells isolated from stromal vascular fraction (SVF) of human adipose tissue samples. (A)** SVF cells were stained with fluorescent antibodies and mesenchymal and hematopoietic subpopulations were purified by FACS using the representative gating strategy shown. Debris was excluded based on the ratio of forward scatter (FSC) height to side scatter (SSC) height. Clusters and aggregates were excluded based on the ratio of SSC width to SSC height. Live cells were DAPI negative. CD45 fluorescence revealed distinct positive and negative populations. The CD45NEG population represented mesenchymal lineage cells, which included CD31NEG/CD29POS conventional mesenchymal adipocyte precursor cells. The CD45POS population contained multiple populations of hematopoietic lineage cells: red gate - myeloid lineage (CD14POS), including purple gate - macrophages (ATM), yellow gate – monocytes, and blue gate – neutrophils; orange gate – basophils; green gate – lymphocytes (CD14NEG). **(B and C)** Representative images for identification of hematopoietic lineage subpopulations by flow cytometry from two different cell donors. The flow cytometry strategy was identical to the first four panels in A, with addition of fluorescent antibodies for the identification of hematopoietic subpopulations. **(B)** Basophils: CD14NEG/Siglec-8POS; Neutrophils: CD14POS, CD16POS, HLA-DRNEG, Siglec-8NEG. **(C)** ATM and monocytes: CD14POS/HLA-DRPOS with divergent expression of CD206 (ATM CD206POS, monocytes CD206NEG). Lymphocytes: CD14NEG/CD206NEG. ATM and monocyte populations were pooled for analysis because of difficulty with complete discrimination and low numbers for downstream analysis. This image and legend has been adapted with author permission from reference [18] in the main text.

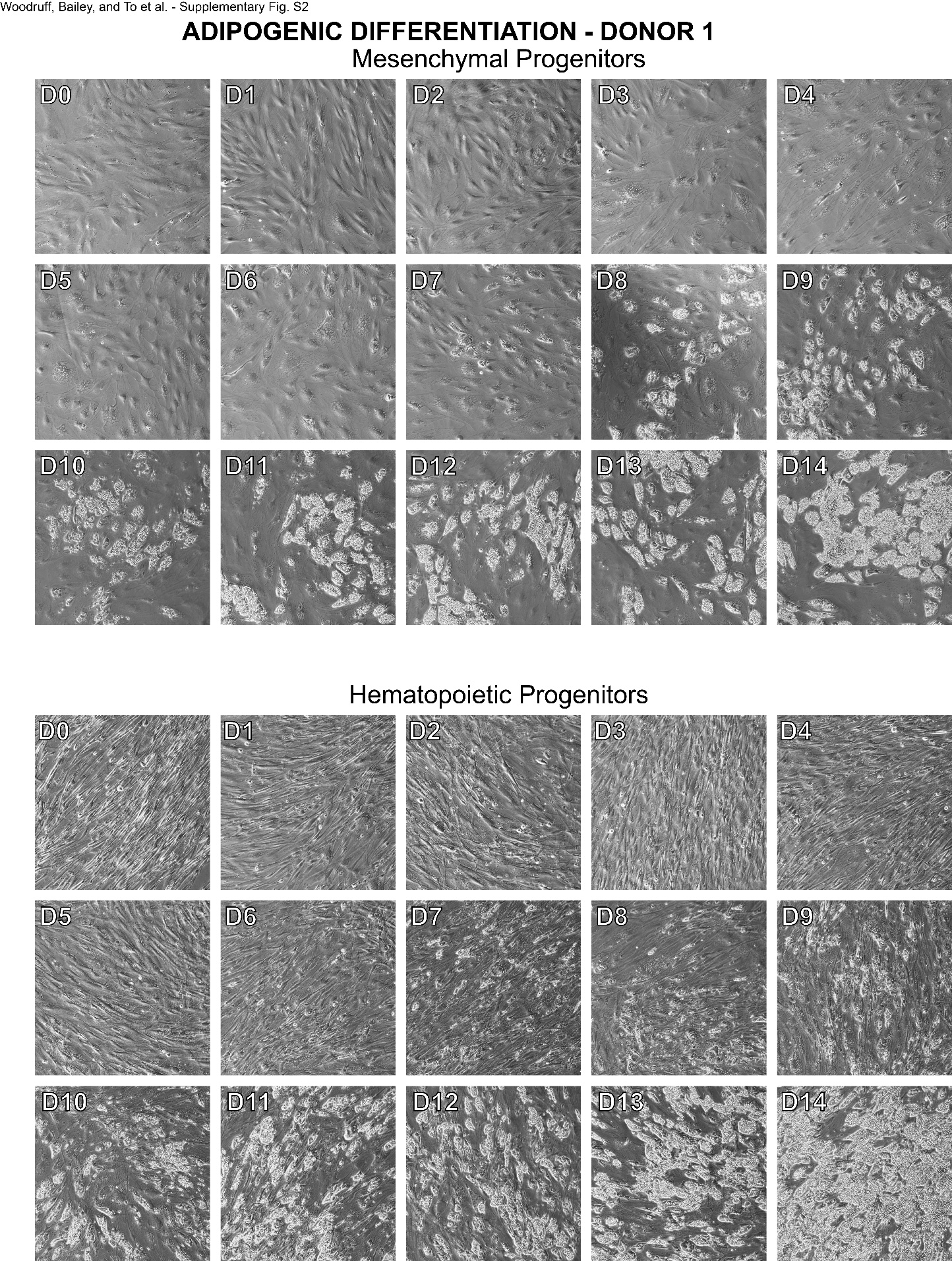

**Fig. S2. Adipogenic differentiation of mesenchymal and hematopoietic progenitor cells from human female Donor 1.** Brightfield images of cultured cells were taken daily, starting with a confluent culture of undifferentiated cells (D0) in Growth Media (GM). Differentiation was started on D1 with the addition of Complete Differentiation Media (CDM), which was changed every two days until D8, at which point media was exchanged for Adipocyte Maintenance Media (AMM). AMM was changed every two days until D14. Media recipes are given in Materials and Methods.

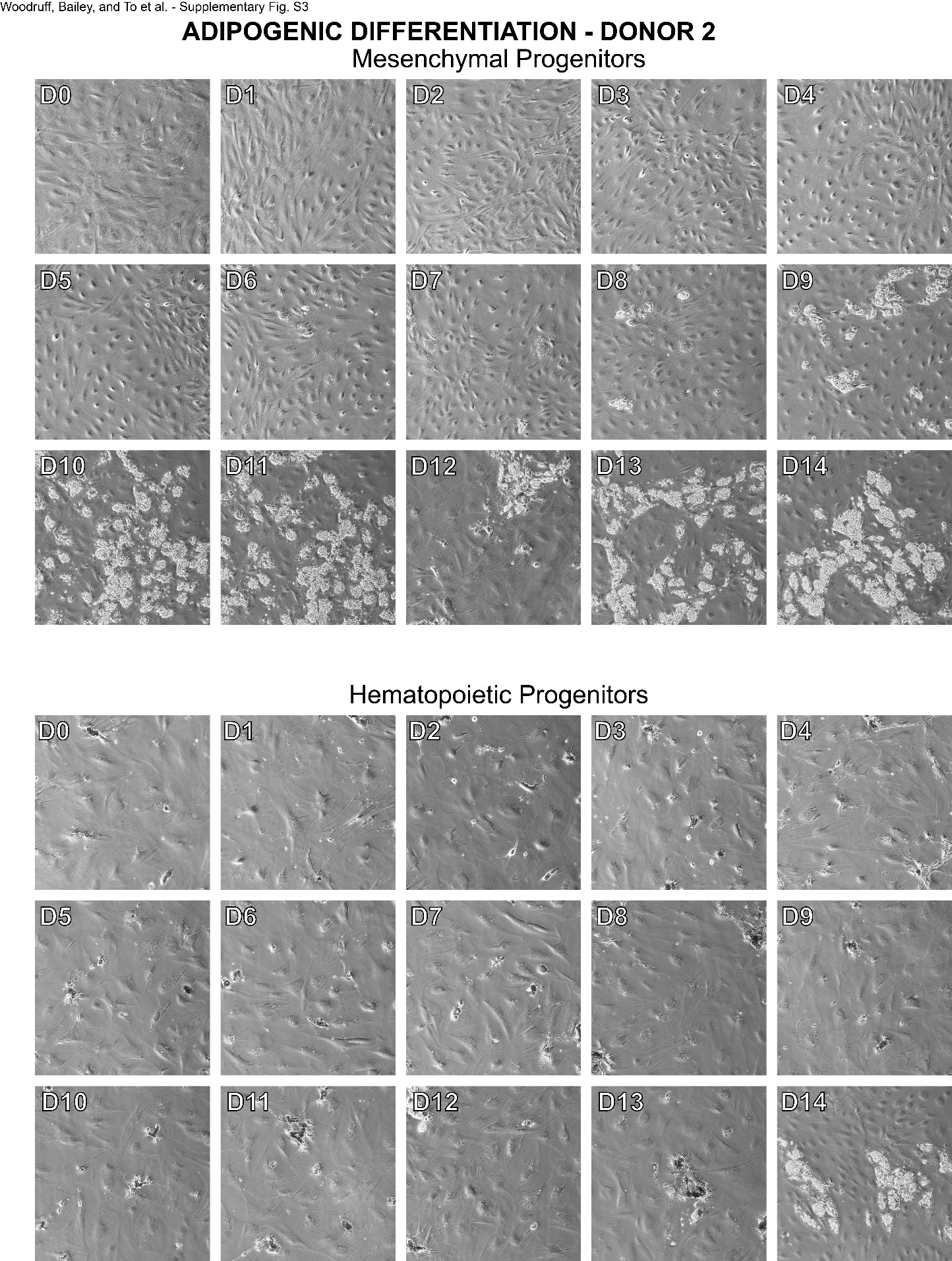

**Fig. S3. Adipogenic differentiation of mesenchymal and hematopoietic progenitor cells from human female Donor 2.** Brightfield images of cultured cells were taken daily, starting with a confluent culture of undifferentiated cells (D0) in Growth Media (GM). Differentiation was started on D1 with the addition of Complete Differentiation Media (CDM), which was changed every two days until D8, at which point media was exchanged for Adipocyte Maintenance Media (AMM). AMM was changed every two days until D14. Media recipes are given in Materials and Methods.

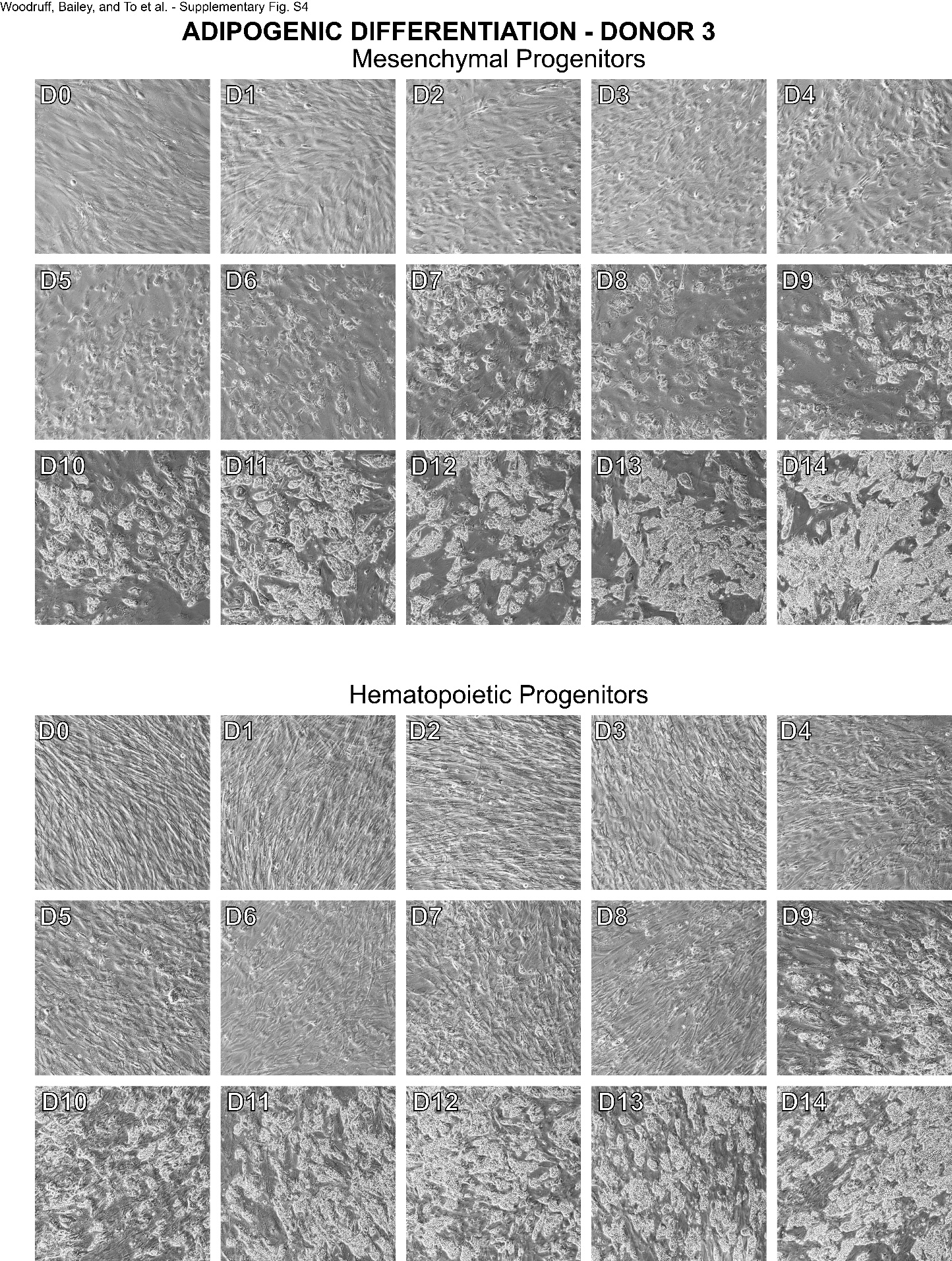

**Fig. S4. Adipogenic differentiation of mesenchymal and hematopoietic progenitor cells from human female Donor 3.** Brightfield images of cultured cells were taken daily, starting with a confluent culture of undifferentiated cells (D0) in Growth Media (GM). Differentiation was started on D1 with the addition of Complete Differentiation Media (CDM), which was changed every two days until D8, at which point media was exchanged for Adipocyte Maintenance Media (AMM). AMM was changed every two days until D14. Media recipes are given in Materials and Methods.

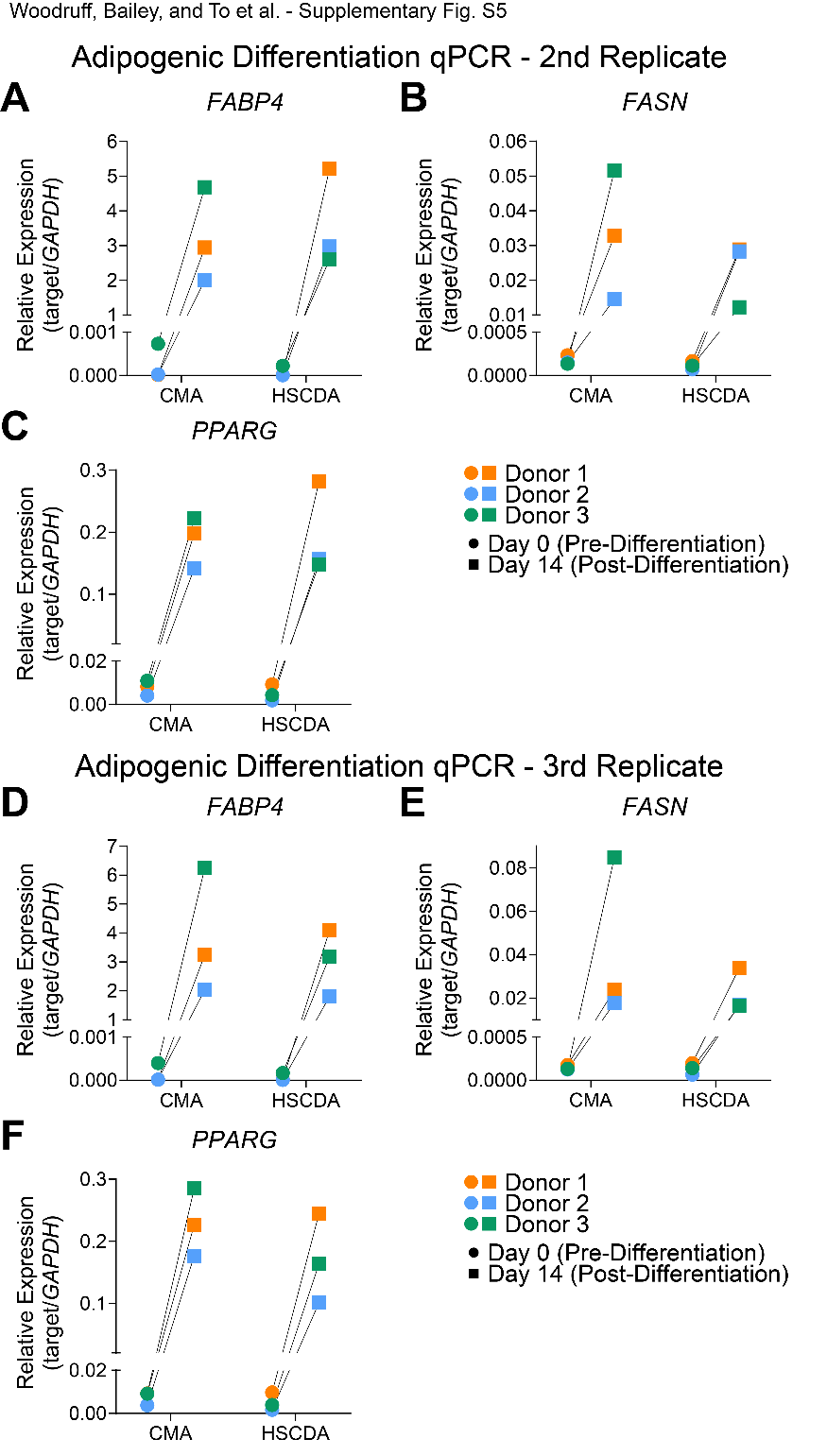

**Fig. S5. Human hematopoietic and mesenchymal precursors undergo similar adipogenic differentiation *in vitro*. (A-C)** Second replicate experiment. Confluent human precursor cells and fully differentiated HSCDAs and CMAs were analyzed by RT-qPCR for mRNA expression of *FABP4*, *FASN*, and *PPARG*. Each data point is the mean of triplicate reactions, and expression is normalized to *GAPDH*. Paired data points show relative expression for cells from each donor at D0 (circles) and D14 (squares). **(D-F)** Third replicate experiment. Same as **A-C**.

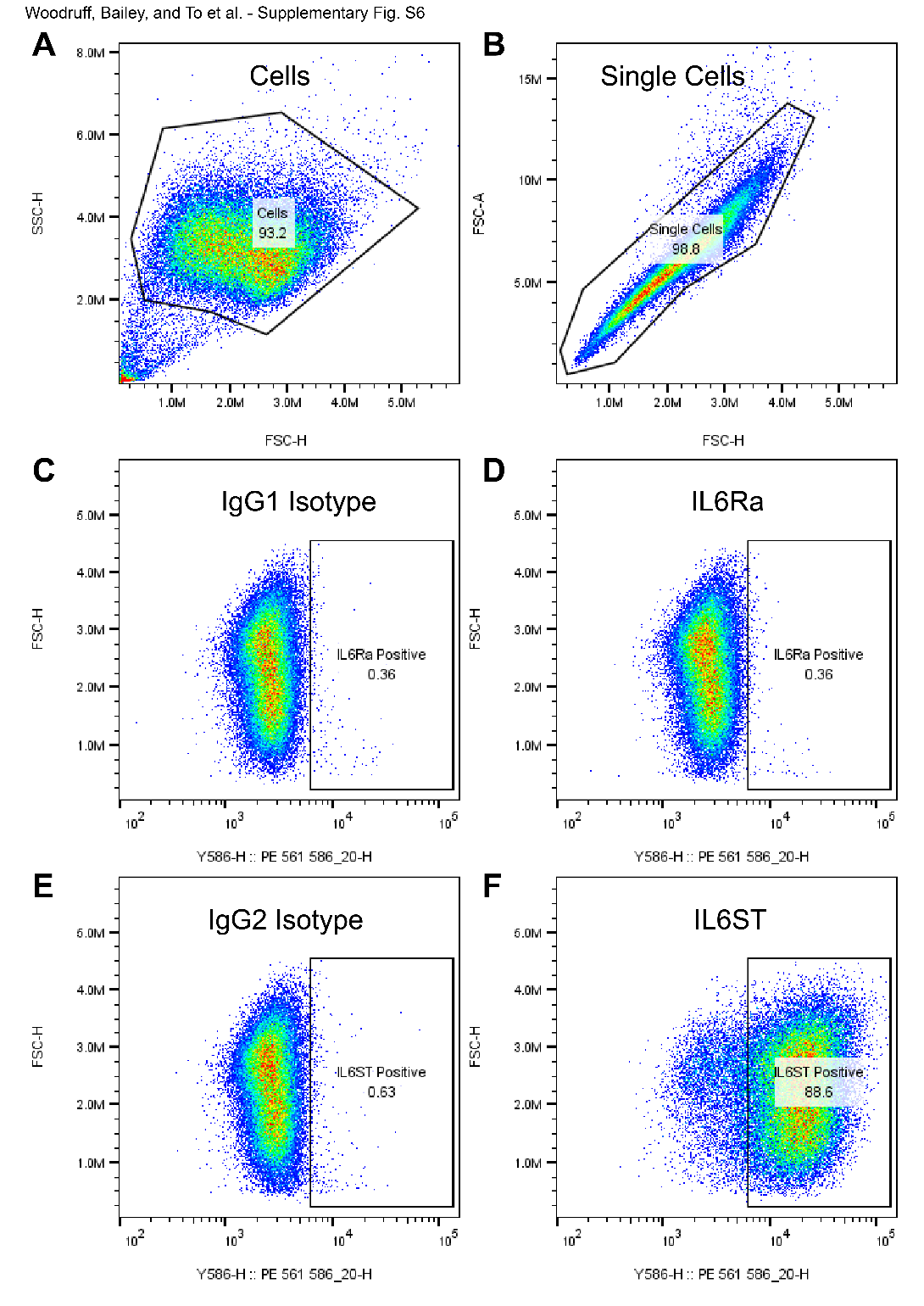

**Fig. S6. Flow gating for PEO1 cell surface expression of IL6Ra and IL6ST.** **(A)** Unstained PEO1 cells were gated to include cells but exclude debris and subsequently gated for **(B)** single cells. These gates were then applied to cells stained with **(C)** PE-conjugated IgG1 isotype, which was used to set the positive gate for **(D)** IL6Ra, or to cells stained with **(E)** PE-conjugated IgG2 isotype, which was used to set the positive gate for **(F)** IL6ST.

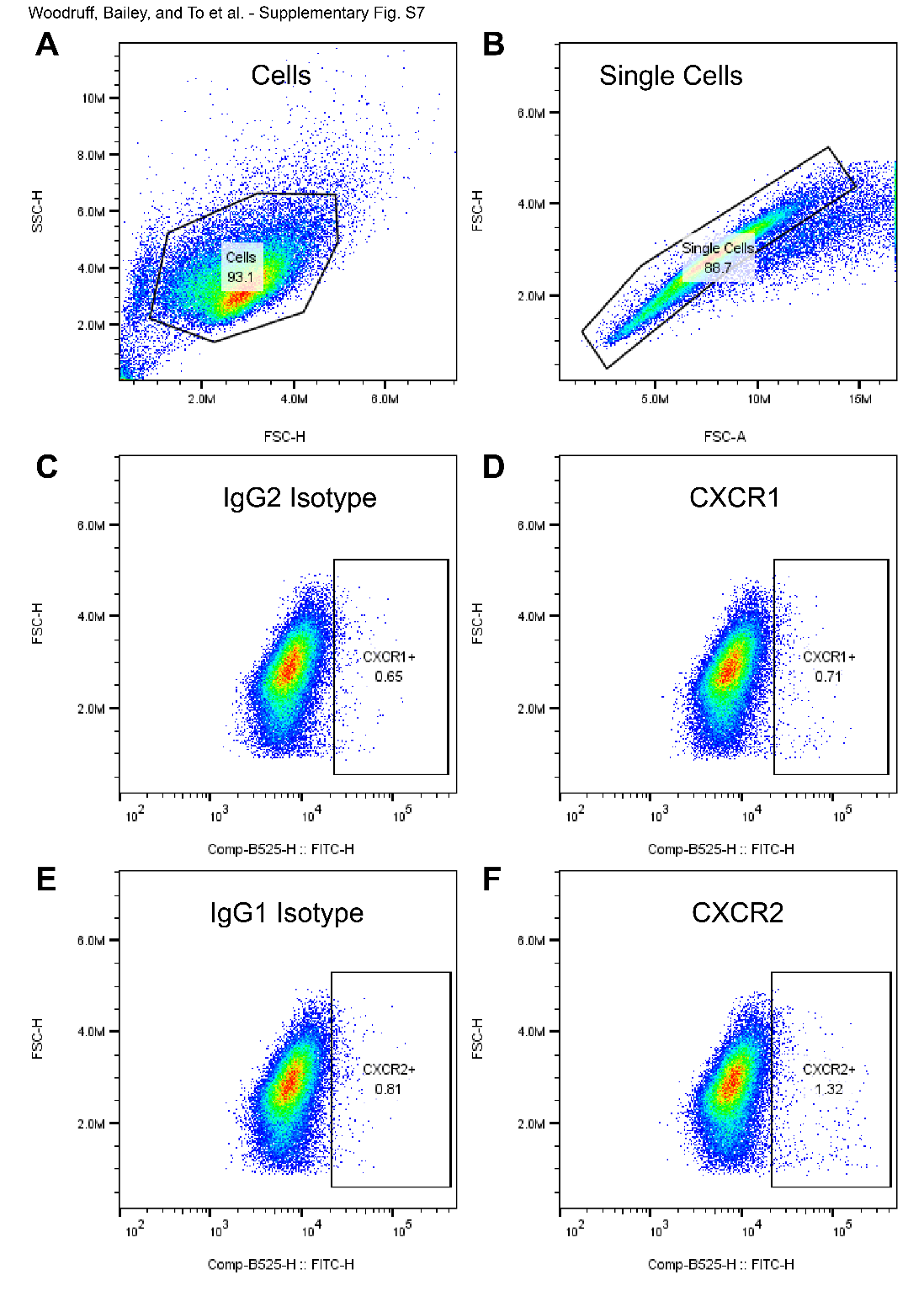

**Fig. S7. Flow gating for PEO1 cell surface expression of CXCR1 and CXCR2.** **(A)** Unstained PEO1 cells were gated to include cells but exclude debris and subsequently gated for **(B)** single cells. These gates were then applied to cells stained with **(C)** PE-conjugated IgG2 isotype, which was used to set the positive gate for **(D)** CXCR1, or to cells stained with **(E)** PE-conjugated IgG1 isotype, which was used to set the positive gate for **(F)** CXCR2.

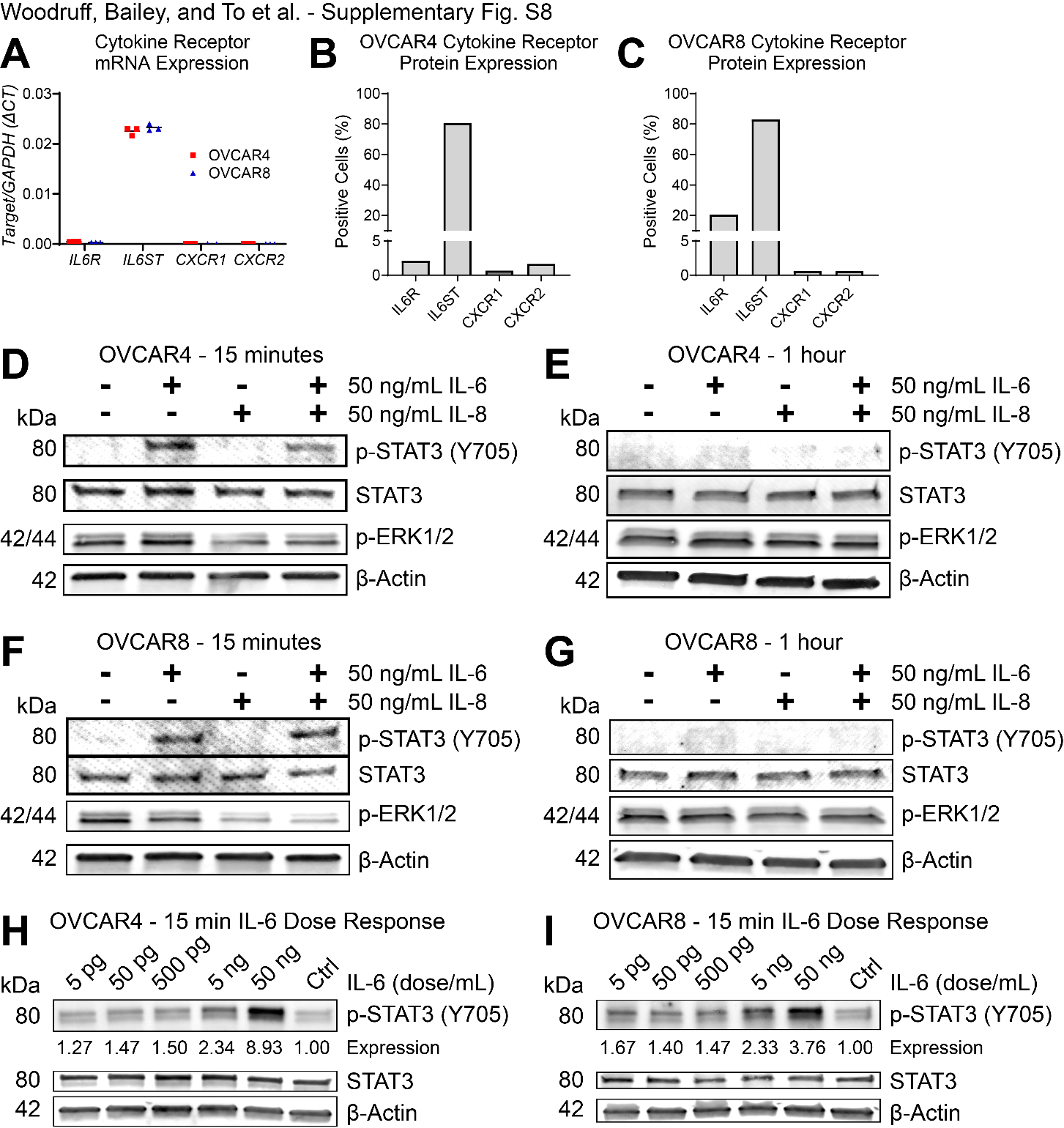

**Fig. S8. HGSC Cell Cytokine Receptor Expression.** **(A)** RNA from OVCAR4 and OVCAR8 HGSC cells was assayed by RT-qPCR for the indicated cytokine receptor transcripts. *GAPDH* was used for ΔCt quantification. **(B-C)** OVCAR4 and OVCAR8 cells were assayed by flow cytometry for surface protein expression of the indicated cytokine receptors. **(D-I)** OVCAR4 and OVCAR8 cells were incubated in media supplemented with the indicated cytokines. Protein lysates were assayed by immunoblot for total and phosphorylated STAT3 (IL-6 signaling), phosphorylated ERK1/2 (IL-8 signaling) and β-actin loading control. For **H-I**, p-STAT3 was quantified by densitometry and normalized to control (no added IL-6).

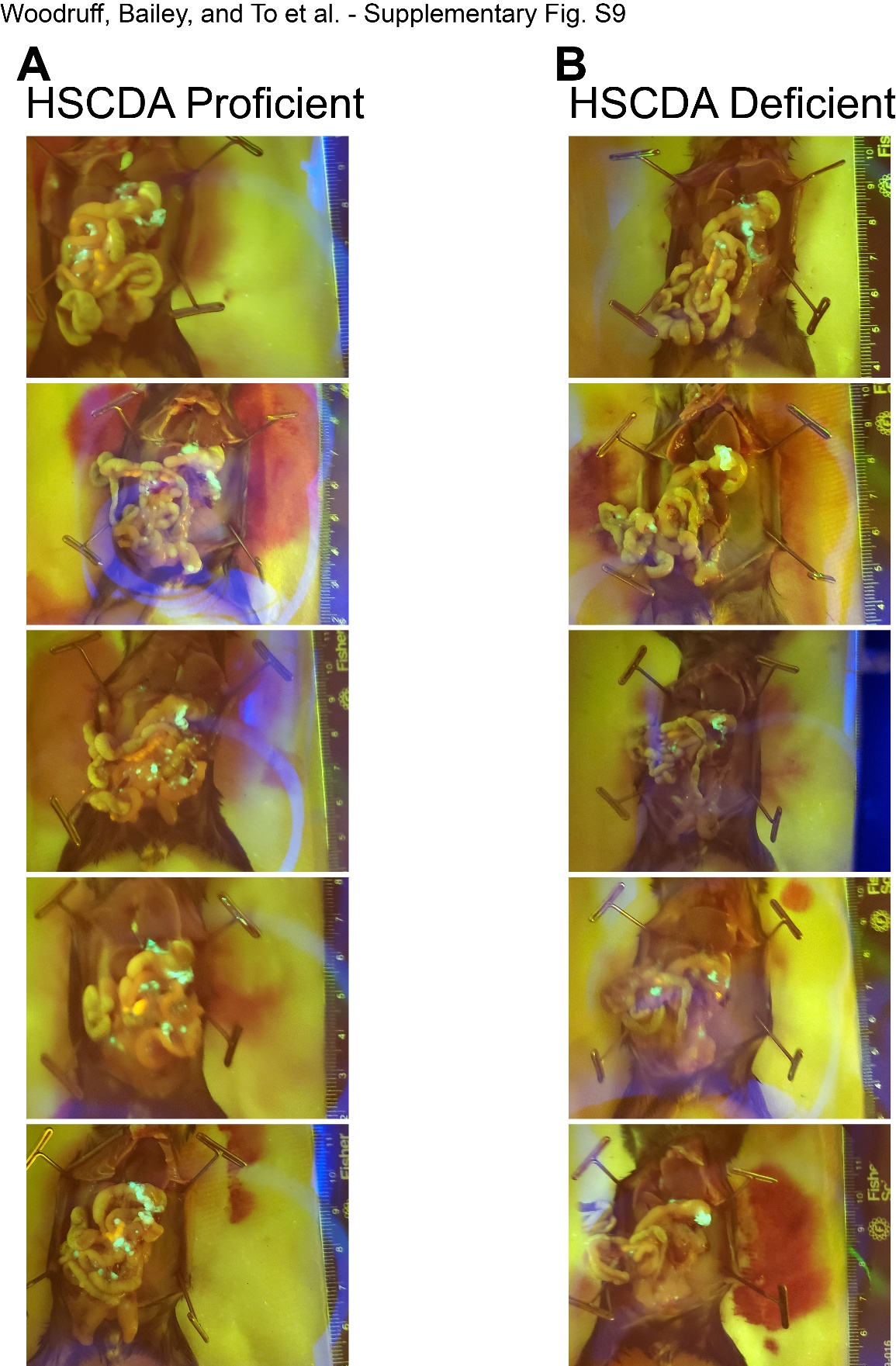

**Fig S9. Representative GFP necropsy images of ID8 omentum and disseminated tumors in HSCDA Proficient and Deficient mice. (A)** HSCDA Proficient. **(B)** HSCDA Deficient. 39 days after cell injection, mice were euthanized and necropsy was performed. ID8 cells are GFP+. Omentum and disseminated tumors are visible within the open peritoneum.

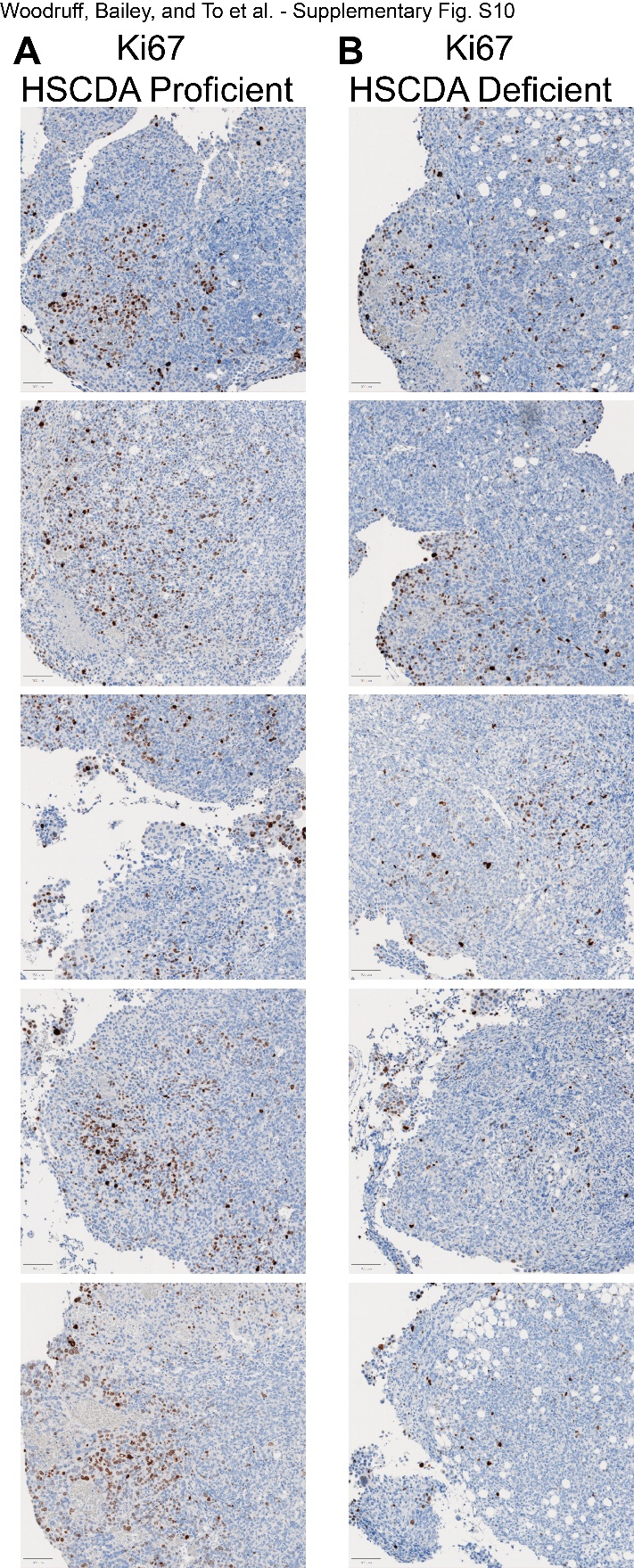

**Fig. S10. Representative brightfield IHC images of Ki67 staining in ID8 omentum tumors from HSCDA Proficient and Deficient mice. (A-B)** Ki67 IHC stained ID8 omentum tumors. Scale bars (bottom left of each image) = 100 µm.

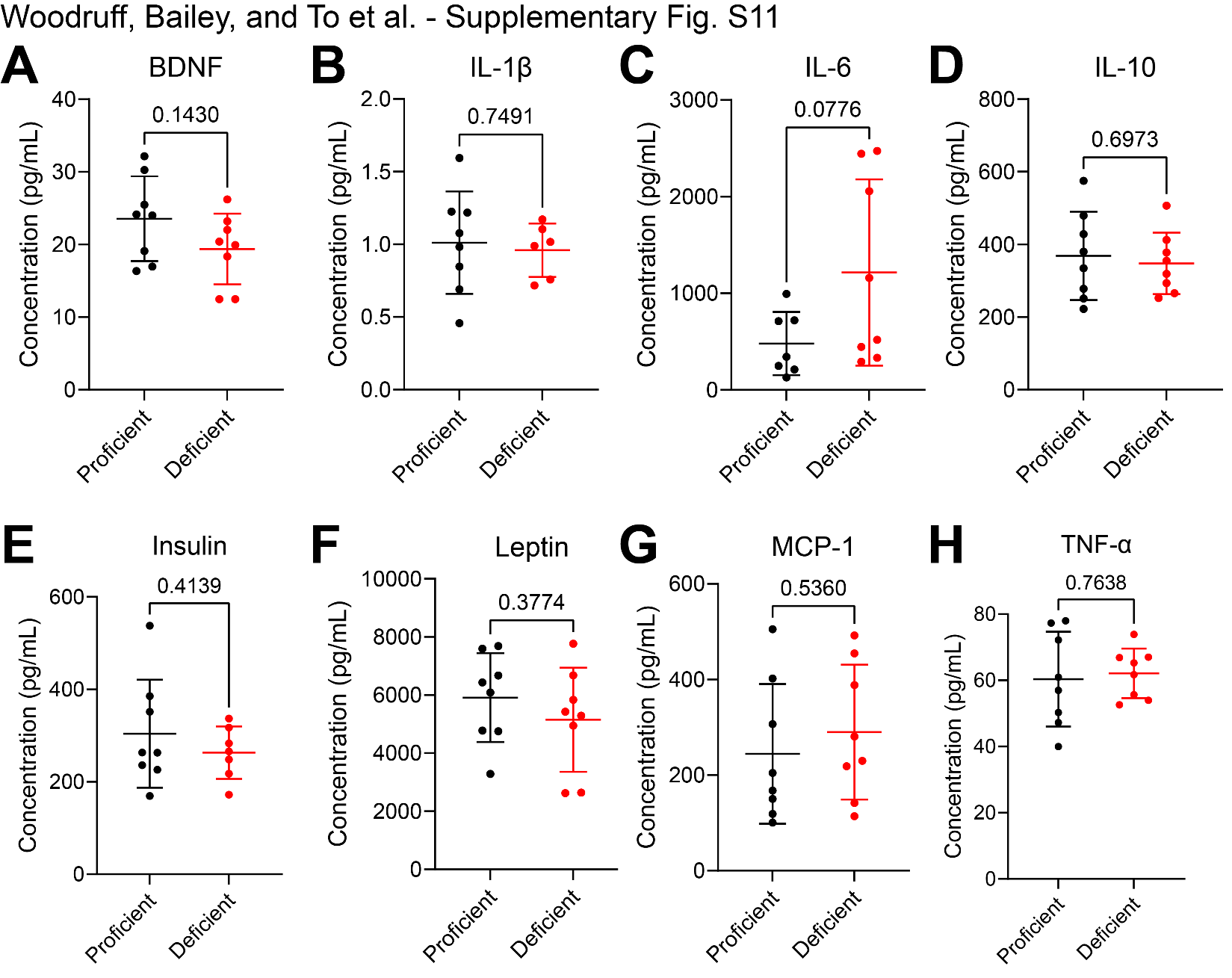

**Fig. S11. Cytokine milieu within ascites fluid of ID8 tumor-bearing mice.** Ascites fluid from tumor-bearing mice was collected. Fluid was cleared by centrifugation, and cytokine concentration was assayed in the supernatant by ELISA. Data are plotted as individual data points with overlaid mean ± SD. Unpaired t-test.

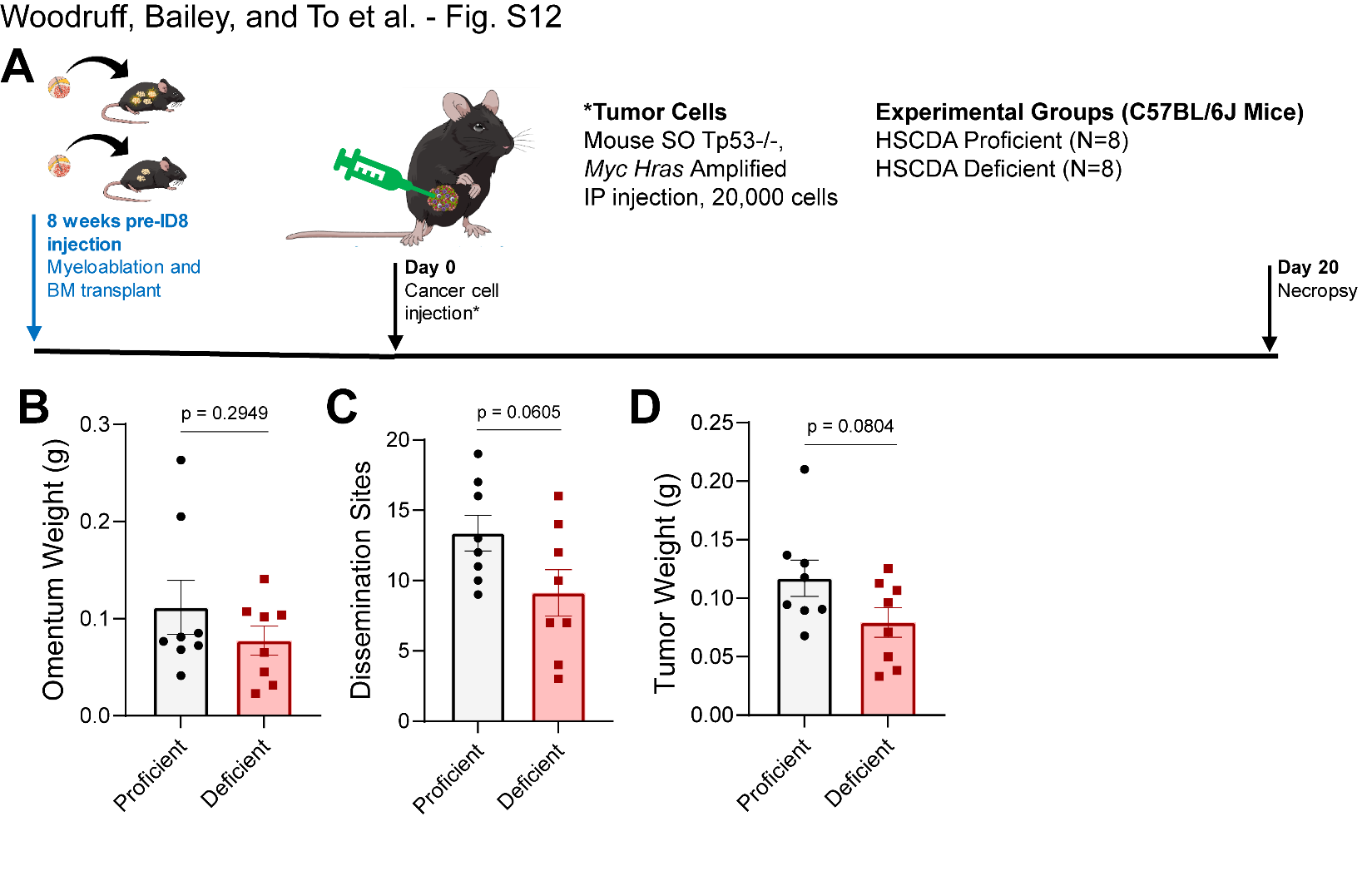

**Fig. S12. SO model intraperitoneal tumor burden is modestly lower in HSCDA Deficient mice. (A)** Experimental timeline, including bone marrow transplant and subsequent tumor study. **(B)** Weight of omentum tumors. **(C)** Number of disseminated tumor nodules. **(D)** Weight of disseminated tumor nodules. For **(B-D)**, graphs show individual data points with mean ± SEM. Unpaired t-test.

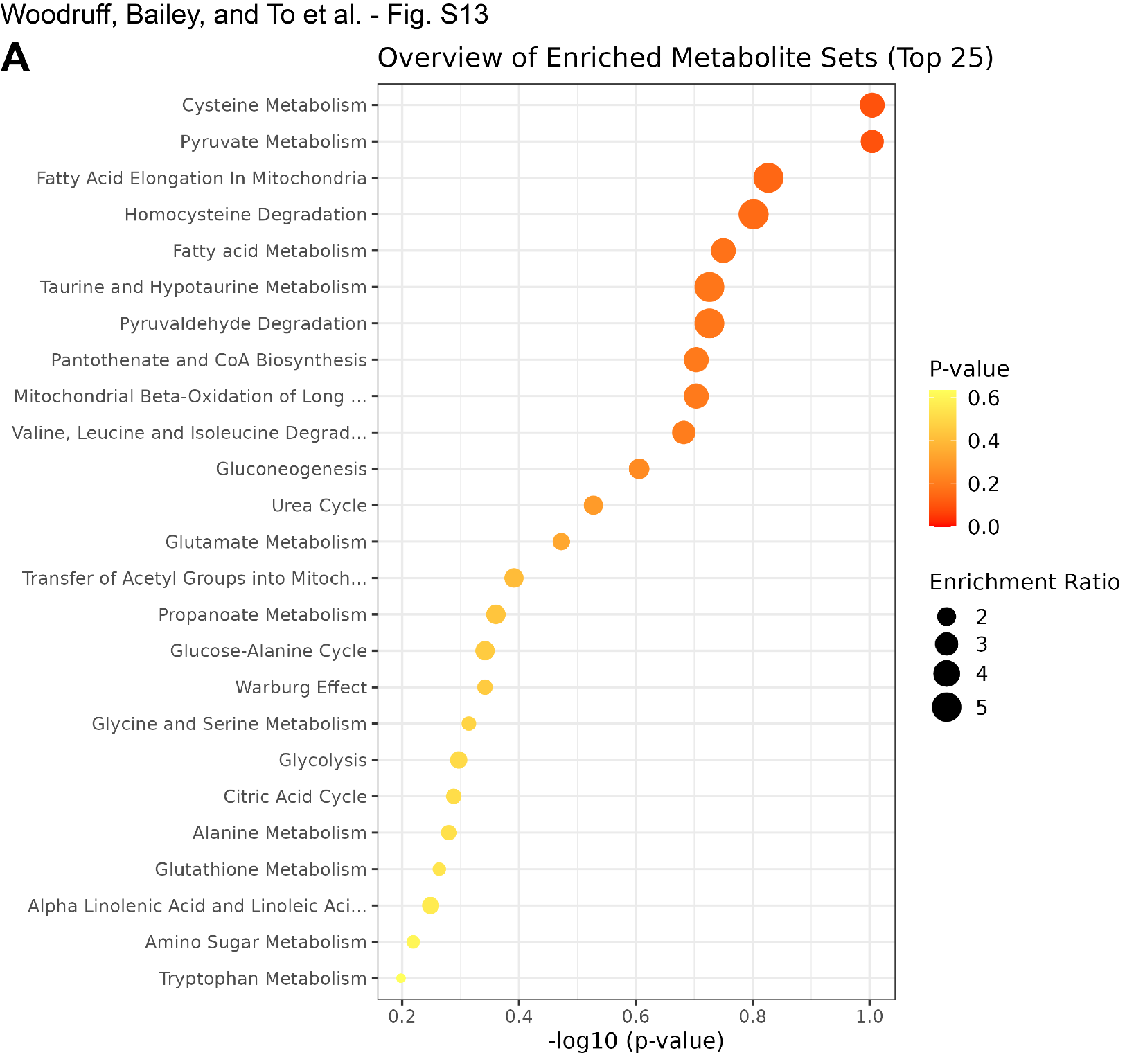

**Fig. S13. Metabolic pathways are differentially regulated in human HGSC ascites in an IL-6-depenent manner. (A)** Cell-free human HGSC ascites were analyzed by mass spec for lipid and metabolic content. Samples were divided into IL-6 low (n=10) and IL-6 high (n=10) and Overrepresentation Analysis (ORA) was performed to predict differentially regulated metabolic pathways. The original dataset is published at <https://doi.org/10.1158/1078-0432.CCR-20-1762> (reference #24 in the main text).

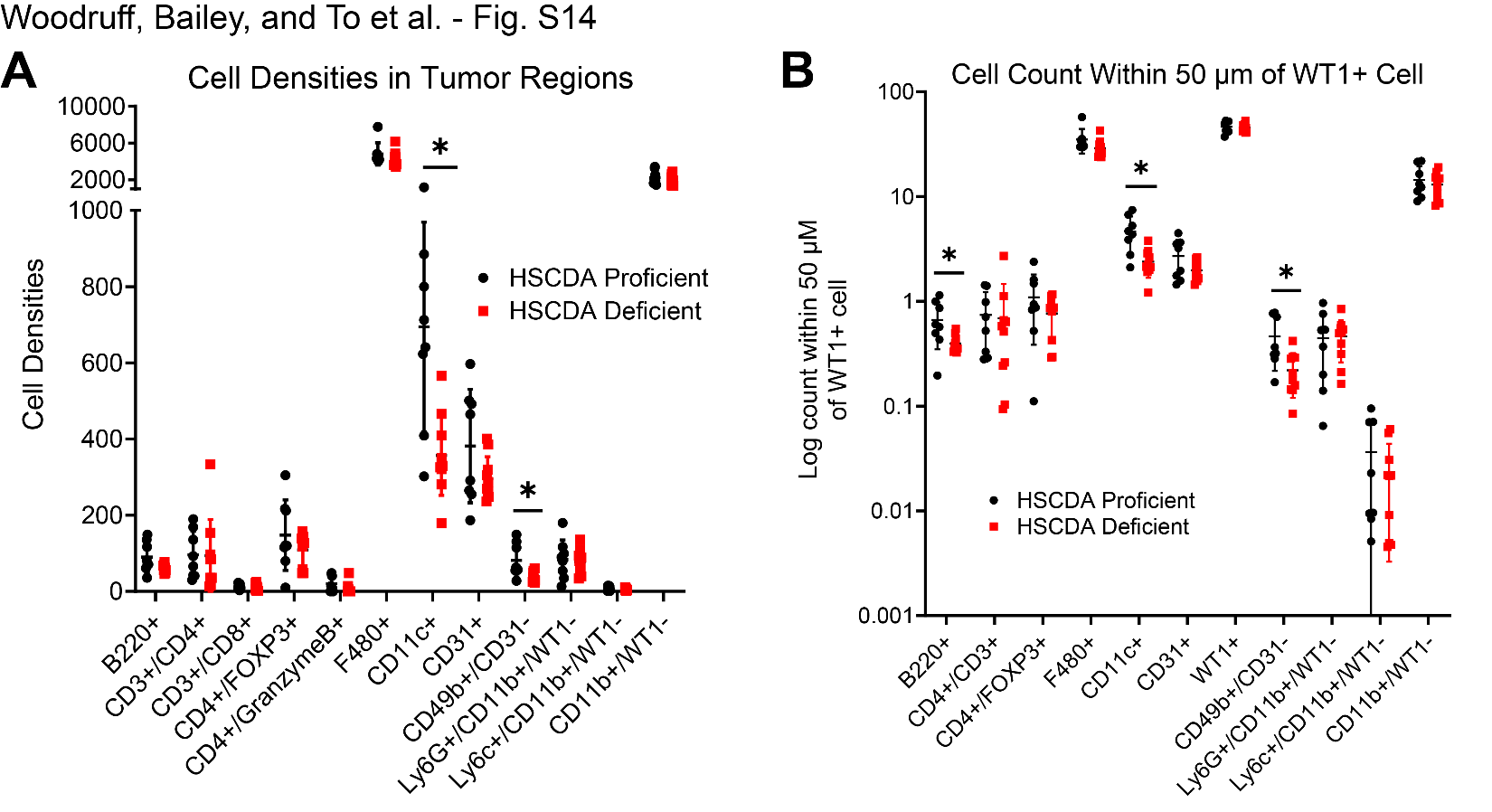

**Fig. S14. HSCDA ablation reduces dendritic cell and natural killer cell density in ID8 omentum tumors.** Omentum tumors from HSCDA Proficient and Deficient mice were stained for the indicated immune cell markers. **(A)** Quantification of cells per mm^2^ of tissue in omentum tumors. **(B)** Number of cells within 50 microns of a WT1+ tumor cell. Error bars = SD. Unpaired t-test. *p<0.05.

**Supplementary Table ST1. RT-qPCR Primers**

| **Name** | **Sequence** | **Target** | **Amplicon Size** |
| --- | --- | --- | --- |
| GAPDH-F | GTCTCCTCTGACTTCAACAGCG | Human *GAPDH* | 131 bp |
| GAPDH-R | ACCACCCTGTTGCTGTAGCCAA |  |  |
| IL6R-F | AGTGTCGGGAGCAAGTTCAG | Human *IL6R* | 232 bp |
| IL6R-R | GGAGGTCCTTGACCATCCAT |  |  |
| IL6ST-F | GGGTAGAAGCAGAGAATGCCC | Human *IL6ST* | 243 bp |
| IL6ST-R | CTGTGTCTTCAGGAGGAATCTGG |  |  |
| CXCR1-F | CAGCTCCTACTGTTGGACACA | Human *CXCR1* | 221 bp |
| CXCR1-R | GAACACTAGGGCATAGGCGA |  |  |
| CXCR2-F | CACTCCAATAACAGCAGGTCACA | Human *CXCR2* | 211 bp |
| CXCR2-R | ACATGGGGCGGCATCTAGTA |  |  |
| FABP4-F | ACAGGAAAGTCAAGAGCACCAT | Human *FAPB4* | 152 bp |
| FABP4-R | AACTCTCGTGGAAGTGACGC |  |  |
| FASN-F | TCGTGTTGACTTCTCGCTCC | Human *FASN* | 199 bp |
| FASN-R | CCATCTCTCAAGACCACGGC |  |  |
| PPARG-F | GTGAAGGATGCAAGGGTTTCT | Human *PPARG* | 300 bp |
| PPARG-R | GCGGGAAGGACTTTATGTATGAG |  |  |

**Supplementary Table ST2. Flow Cytometry Antibodies**

| **Target** | **Fluorophore** | **Concentration** | **Supplier** | **Catalog #** | **RRID** |
| --- | --- | --- | --- | --- | --- |
| Human IL6R | PE | 1:100 | BioLegend | 352804 | AB_10900066 |
| IgG1 Isotype | PE | 1:100 | BioLegend | 400114 | N/A |
| Human IL6ST | PE | 1:100 | BioLegend | 362004 | AB_2563402 |
| IgG2 Isotype | PE | 1:100 | BioLegend | 400214 | N/A |
| Human CXCR1 | FITC | 1:100 | BioLegend | 320606 | AB_439813 |
| IgG2 Isotype | FITC | 1:100 | BioLegend | 400310 | N/A |
| Human CXCR2 | FITC | 1:100 | BioLegend | 320704 | AB_439805 |
| IgG1 Isotype | FITC | 1:100 | BioLegend | 400110 | N/A |

**Supplementary Table ST3. Immunoblot Primary Antibodies**

| **Target** | **Host Species** | **Concentration** | **Supplier** | **Catalog #** | **RRID** |
| --- | --- | --- | --- | --- | --- |
| p-STAT3* | Rb | 1:500 | Cell Signaling | 9145 | AB_2491009 |
| Total STAT3 | Ms | 1:500 | Cell Signaling | 9139 | AB_331757 |
| p-ERK1/2 | Rb | 1:1000 | Cell Signaling | 4695 | AB_390779 |
| Β-Actin | Ms | 1:10000 | Abcam | ab6276 | AB_2223210 |

**Supplementary Table ST4. Immunoblot Secondary Antibodies**

| **Name** | **IR Dye** | **Concentration** | **Supplier** | **Catalog #** | **RRID** |
| --- | --- | --- | --- | --- | --- |
| Goat anti-rabbit* | 800CW | 1:20000 | LI-COR | 926-32211 | AB_621843 |
| Goat anti-mouse | 800CW | 1:20000 | LI-COR | 925-32210 | AB_2687825 |
| Goat anti-mouse | 680RD | 1:20000 | LI-COR | 926-68070 | AB_10956588 |

*****For sensitive quantification of p-STAT3 in low dose IL-6 experiments **(Fig. 3 and S8)**, we used HRP-linked goat anti-rabbit IgG secondary antibody (Cell Signaling 7074, RRID: AB_2099233) at 1:5000 followed by detection with SuperSignal West Femto ECL substrate (Thermo 34095).

**Supplementary Table ST5. Multiplex IHC Antibodies**

| **Target** | **Supplier** | **Catalog #** | **Conc.** | **RRID** | **OPAL** | **OPAL Conc.** |
| --- | --- | --- | --- | --- | --- | --- |
| Ms CD11b | Novus | 89474 | 1:1000 | AB_1216361 | 480 DI | 1:150 |
| Ms CD11c | Cell Signaling | 97585 | 1:100 | AB_2800282 | 520 DI | 1:150 |
| Ms Ly6c | BioRad | MCA2389 | 1:250 | AB_844552 | 540 DI | 1:150 |
| Ms Ly6g | Cell Signaling | 87048 | 1:100 | AB_2909808 | 620 DI | 1:150 |
| Ms CD49b | Invitrogen | MA5-32306 | 1:50 | AB_2809588 | 650 DI | 1:150 |
| Ms CD31 | Abcam | Ab182981 | 1:100 | AB_2920881 | 690 DI | 1:150 |
| Ms F480 | Cell Signaling | 30325 | 1:100 | AB_2798990 | 780 | 1:25 |
| Ms CD3 | Cell Signaling | 99940 | 1:50 | AB_2755035 | 540 DI | 1:150 |
| Ms CD4 | Invitrogen | 14-9766-82 | 1:200 | AB_2573008 | 650 DI | 1:150 |
| Ms CD8 | Cell Signaling | 98941 | 1:400 | AB_2756376 | 480 DI | 1:150 |
| Ms Granzyme B | Cell Signaling | 44153 | 1:100 | AB_2857976 | 520 DI | 1:150 |
| Ms Foxp3 | R&D Systems | MAB8214 | 1:200 | AB_2929004 | 690 DI | 1:150 |
| Ms B220 | BDPharm | 557390 | 1:1000 | AB_396673 | 620 DI | 1:150 |
| Ms WT1 | Novus | 110-60011 | 1:400 | AB_1849479 | 570 DI | 1:150 |
| Ms CD80 | Invitrogen | PA5-85913 | 1:100 | AB_2802714 | 480 DI | 1:300 |
| Ms MHCII | BioLegend | 107601 | 1:100 | AB_313316 | 650 DI | 1:150 |
| Ms NCR1 | Abcam | ab233558 | 1:100 | AB_2904203 | 520 DI | 1:300 |
| Ms Tbet | Cell Signaling | 97135 | 1:400 | AB_3675945 | 570 DI | 1:100 |
